# G4SCOPE identifies sequence features and epigenetic contexts of chromatin G-quadruplexes

**DOI:** 10.64898/2026.09.28.748631

**Authors:** Armaghan Sarvar, Isabel Esain-Garcia, Larry Melidis, Damian Yap, Ciara O’Flanagan, Viviana Cerda, Daniel Lai, David Tannahill, Yongjin Park, Shankar Balasubramanian, Samuel Aparicio

## Abstract

Cancer-associated epigenetic remodeling involves a complex interplay between DNA secondary structures and covalent cytosine modifications, yet predicting endogenously folded, chromatin-constrained G-quadruplex (G4) dynamics remains challenging. Here, we develop G4SCOPE, a multi-task deep learning framework that models chromatin-associated G4 occupancy and reconstructs quantitative high-resolution CUT&Tag profiles directly from primary DNA sequence alone. We initially trained G4SCOPE on genotype-stratified G4 enriched cut&tag landscapes generated across an isogenic series of human breast epithelial cells (WT, TP53−/−, TP53−/−;BRCA1−/−, and TP53−/−;BRCA2−/−). G4SCOPE integrates convolutional motif learning with transformer self-attention to decode long-range sequence contexts. The model robustly distinguishes chromatin-associated G4s from GC-matched loci with intrinsic in vitro folding potential, generalizes across distinct breast lineages, and transfers accurately to an independent K562 cellular context. Strikingly, genotype-resolved mapping reveals a profound divergence in genome maintenance states: BRCA1 deficiency indicated a global inflation of chromatin-embedded G4 density, whereas BRCA2 loss was associated with systemic, chromosome-wide structural depletion. Furthermore, unsupervised clustering of model-derived latent sequence representations identified distinct biological G4 subclasses that occupy unique regulatory class genomic environments associated with distinct local 5-methylcytosine and 5-hydroxymethylcytosine landscapes. Together, our framework decodes the primary sequence grammar governing cell-based G4 occupancy and demonstrates that sequence-defined G4 topologies harbor intrinsic and distinct local epigenetic features.

## Introduction

Cancer-associated states are characterized by pervasive changes in epigenetic regulation, including both covalent cytosine modifications and DNA secondary structures that shape chromatin organization and transcription^1–3^. Among these, G-quadruplexes (G4s) are especially relevant, because, although encoded by DNA sequence, their formation in cells is strongly influenced by chromatin context.

G-quadruplexes are non-canonical nucleic acid secondary structures formed by guanine-rich DNA or RNA sequences that fold into four-stranded assemblies of stacked guanine tetrads stabilized by Hoogsteen hydrogen bonding and monovalent cations^4–6^. Early biochemical and structural studies established that G4s can fold under near-physiological conditions in oligonucleotide sequences found in the genome^4,7,8^. Although G4-forming potential is often captured by canonical motifs composed of four G-tracts separated by loops, G4s can also arise from non-canonical sequence patterns such as bulged motifs with interrupted G-tracts, expanding the sequence space beyond simple consensus rules^4– 7,9,10^. Functionally, G4s influence gene regulation, chromatin organization and genome stability rather than acting as passive sequence motifs^11–16^. Because guanine-rich sequences become transiently single-stranded during transcription and replication, folded G4s are associated with regulatory chromatin and nucleosome positioning, and can impede polymerase progression and promote replication stress and DNA damage, particularly when G4-processing pathways are compromised^16–18^. These effects are especially relevant in genotypes with disrupted genome maintenance, including TP53 loss and BRCA1 or BRCA2 deficiency, where altered DNA damage responses, replication-stress tolerance and homologous recombination may change how cells accommodate G4-associated genomic stress^19^.

While both BRCA1 and BRCA2 are foundational to homologous recombination and replication fork protection, they display distinct biochemical roles in managing topological and transcriptional stress. BRCA1 acts as a scaffolding hub orchestrating cell-cycle checkpoints, R-loop resolution, and upstream double-strand break resection. In contrast, BRCA2 functions directly downstream, orchestrating the nucleofilament loading of RAD51 onto single-stranded DNA to secure stalled or reversed replication forks. Prior literature has established a robust synthetic lethality between G4-stabilizing compounds (such as pyridostatin)^20,21^ and general HR deficiency. However, it remains entirely unknown how the complete loss of either BRCA1 or BRCA2 shifts the basal, homeostatic positioning of endogenously folded G4 structures within living chromatin. Whether these genetic backgrounds passively accumulate structural barriers or actively remodel their structural epigenome to maintain survival under chronic replication stress remains a critical, unresolved question in precision oncology.

Genome-wide mapping studies directly illustrate this context dependency. G4-seq maps genome-wide G4-forming sequences in vitro by exploiting polymerase stalling under G4-permissive conditions, providing a comprehensive view of intrinsic folding potential^7,22^. In contrast, cell-based chromatin profiling approaches such as G4 ChIP-seq and G4 CUT&Tag capture endogenous, chromatin-constrained G4 folding and show that only a subset of potential G4-forming loci are actually folded in chromatin, with enrichment at open regulatory regions linked to active transcription^23–27^. Together, these findings show that the presence of a G4- forming sequence does not by itself determine whether the locus is detected as folded in chromatin. For background-set labels throughout this study, we use “in vitro only” to refer to G4-seq loci that were not detected by CUT&Tag in any of the four breast epithelial genotypes.

A similar principle applies to covalent cytosine modifications, including 5-methylcytosine and 5-hydroxymethylcytosine, whose genome-wide distributions are dynamic and context-dependent. Prior work has implicated G4 structures in shaping local DNA methylation landscapes^28^. The key challenge is determining whether folded G4 prediction can also capture epigenetic cytosine-modifications signatures.

Computational approaches to G4 prediction have progressed from motif-based heuristics and sequence scoring frameworks, such as Quadparser, QGRS Mapper, G4Hunter and pqsfinder, to data-driven models trained on high-throughput experimental data^29–33^. Learning-based predictors including Quadron, G4detector, PENGUINN and G4mismatch have improved modeling of intrinsic folding propensity, primarily using G4-seq-derived training data, while DeepG4 and G4Beacon incorporate chromatin accessibility to predict chromatin-associated G4s in cells^34–39^. More recently, hybrid deep-learning architectures have combined convolutional, recurrent and attention-based modules to model sequence features associated with G4 formation^40,41^. However, these methods face several limitations for genotype-stratified, quantitative modeling of endogenous G4 occupancy. They are generally optimized for a single objective, either binary classification or prediction of a stability-related proxy, rather than jointly modeling G4 occupancy and quantitative signal. Many are also trained on in vitro datasets that capture folding propensity rather than chromatin-conditioned state, or depend on auxiliary epigenomic inputs that may not be available across genotypes or conditions^31,37,38^. Notably, although some methods have been trained on G4 ChIP-seq data, to our knowledge, no existing deep learning framework has been developed to predict quantitative G4 CUT&Tag signal intensity, a higher-resolution assay with improved signal-to-noise and reduced background relative to ChIP-seq^24–26,42^.

Most learning-based approaches treat experimentally defined G4 regions as a classification problem rather than predicting the underlying quantitative signal^37,38,43^. We likewise used peak-derived labels for the G4 occupancy task, but jointly trained the model to reconstruct the local quantitative CUT&Tag signal. Because peak definitions can be sensitive to antibody performance, experimental conditions and analytical thresholds, we combined evidence from two independently developed G4-recognizing reagents, BG4 and SG4, and applied stringent reproducibility criteria to define consensus G4-associated loci in each cell line.

The 184-hTERT breast epithelial system provides a controlled, hTERT-immortalized but non-transformed human mammary epithelial background in which genotype-specific effects can be studied without the heterogeneity of cancer-derived lines. The wild-type 184-hTERT-L9 model was established as a clonal line with a normal-like karyotype that retains key features of mammary epithelial biology in culture^44^. This system was subsequently extended to genotype-resolved studies of genome maintenance, first through TP53 loss-of-function branches and later through CRISPRCas9-derived TP53−/−;BRCA1−/− and TP53−/−;BRCA2−/− lineages^45,46^. Here, we generated genome-wide CUT&Tag G4 maps across four isogenic breast epithelial genotypes, WT, TP53−/−, TP53−/−;BRCA1−/− and TP53−/−;BRCA2−/−, and developed G4SCOPE (G4 Sequence-based Chromatin Occupancy and Profile Estimation), a multi-task deep learning framework that models chromatin-associated G4 occupancy directly from DNA sequence (Fig. 1). By training and evaluating the framework across these genotypes, we examined both the shared sequence grammar of G4 occupancy and its performance across genotype-associated G4 landscapes. By jointly predicting G4 occupancy and local CUT&Tag signal, the model uses quantitative assay information as an additional learning objective. This allows the shared sequence encoder to learn features that distinguish G4-positive from G4-negative loci while also capturing finer-scale sequence patterns associated with variation in local signal intensity. The model is therefore trained to identify sequence features linked to chromatin-detected G4 formation rather than intrinsic folding potential alone.

**Figure 1.**
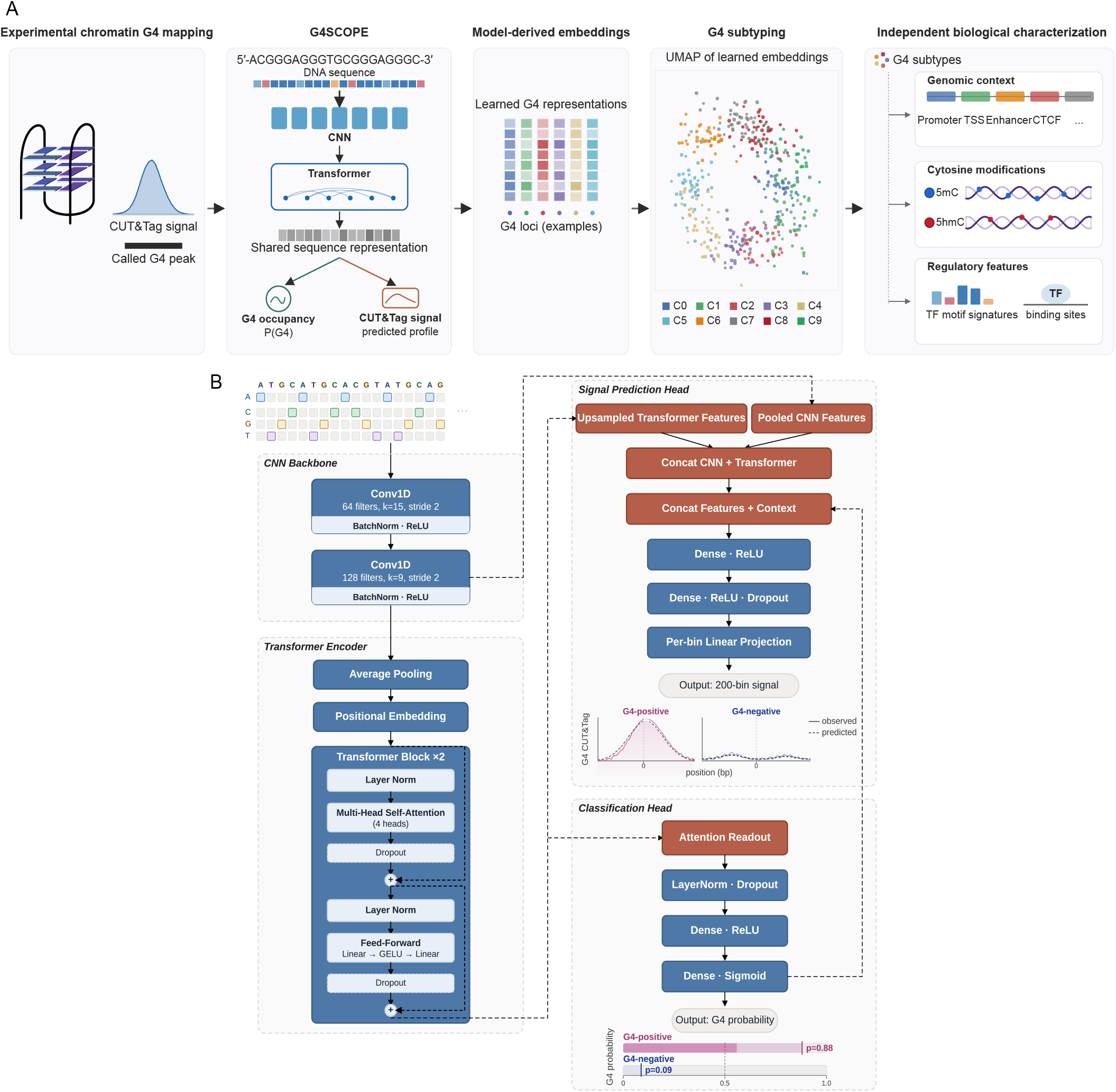
G4SCOPE framework for sequence-based prediction and characterization of chromatin G4s. **(A)** Overview of the study workflow, from experimental chromatin G4 mapping and G4SCOPE modeling to learned representations, G4 subtyping and independent biological characterization. **(B)** G4SCOPE multi-task architecture. One-hot-encoded DNA sequence is processed by a shared CNN–Transformer backbone, followed by parallel heads that predict G4 occupancy and reconstruct the quantitative CUT&Tag signal profile

## Results

### Chromatin G4 landscapes across isogenic breast epithelial genotypes

To define chromatin-associated G4 landscapes across cancer-relevant genotypes, we performed CUT&Tag profiling using the G4-binding reagents BG4 and SG4^47,48^ in an isogenic human mammary epithelial model consisting of immortalized 184-hTERT cells and derivative TP53−/−, TP53−/−;BRCA1−/−, and TP53−/−;BRCA2−/− lines^45^ (see Methods). Hierarchical clustering of genome-wide CUT&Tag profiles showed concordance among biological replicates and genotype-associated structure, with withinreagent Spearman correlations of 0.51–0.80 and BG4–SG4 correlations of 0.36–0.57, across all cell lines, supporting the reproducibility of the assay for each cell line and consistent detection of chromatin-associated G4 structures across them (Fig. 2A).

**Figure 2.**
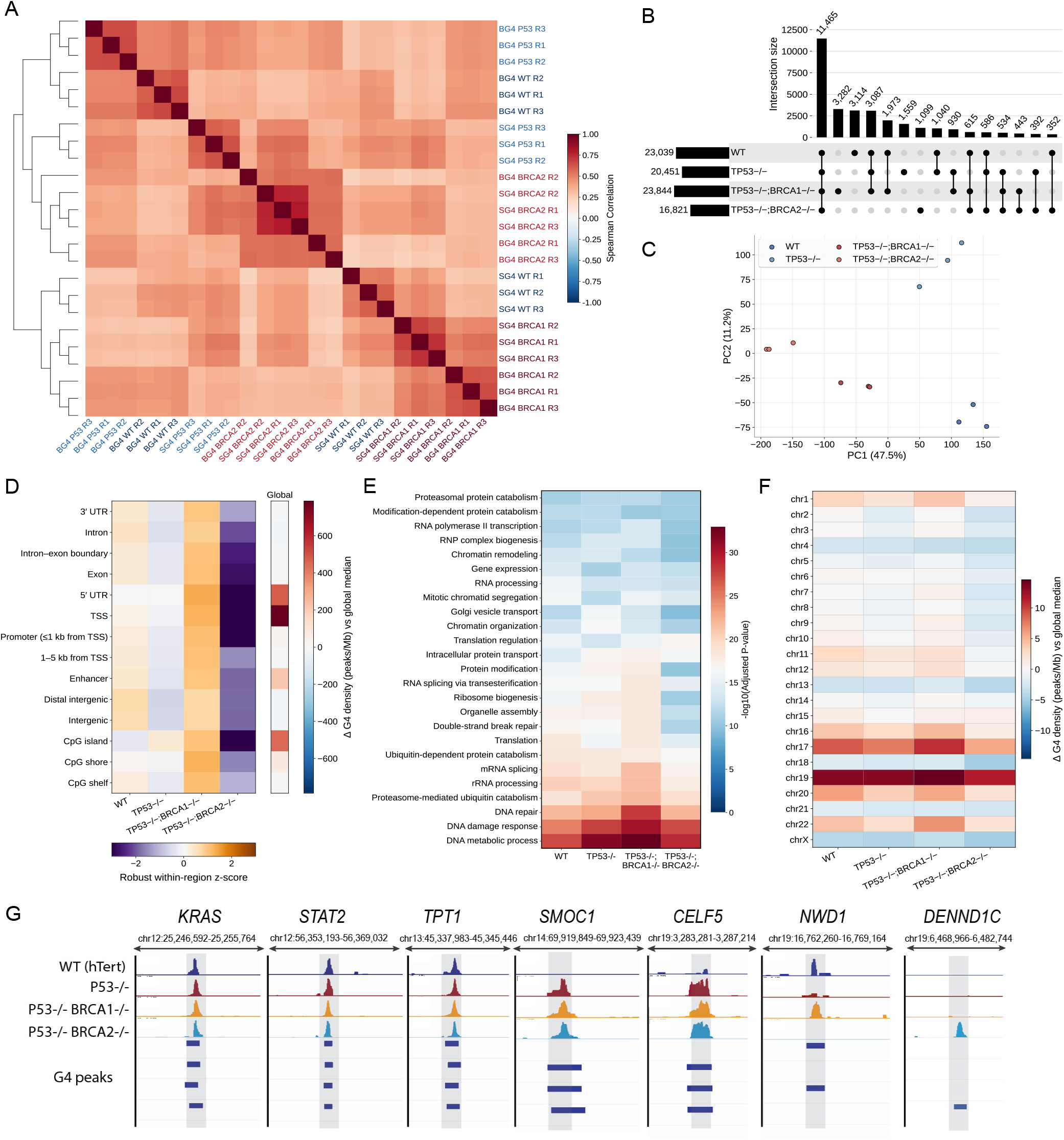
G4 landscapes across isogenic breast epithelial genomes. **(A)** Hierarchical clustering of CUT&Tag signal showing concordance among biological replicates and genotype-associated structure. **(B)** UpSet analysis of 30,471 merged G4 loci showing shared subsets and loci detected in only one genotype. **(C)** Principal component analysis of BG4 CUT&Tag signal showing separation of samples by genotype. **(D)** G4 peak density across genomic annotation classes. Overall density relative to the global median and robust within-annotation z-scores summarize enrichment at promoter-proximal and CpG-associated regions and relative differences among genotypes. **(E)** Functional enrichment of G4-associated regions in each genotype, shown as −log10(adjusted P), highlighting DNA metabolic process, DNA damage response and DNA repair. **(F)** Chromosome-level G4 peak-density differences relative to the global median, showing elevated density on chromosomes 17 and 19, reduced density on chromosome X and lower relative density in TP53−/−;BRCA2−/− cells across most chromosomes. **(G)** Genome-browser tracks at representative loci showing genotype-associated differences in CUT&Tag signal and called peaks.

For each genotype and reagent, replicate-consensus regions were defined as loci detected in at least two of three biological replicates, and genotype-level G4-associated sets were defined by overlap between the BG4 and SG4 consensus regions (Methods). Peak calling using SEACR identified 23,039 G4 loci in WT, 20,451 in TP53−/−, 23,844 in TP53−/−;BRCA1−/−, and 16,821 in TP53−/−;BRCA2−/− cells. Merging CUT&Tag peaks across genotypes yielded a unified set of 30,471 nonredundant loci for direct cross-genotype comparison. Overlap analysis revealed a substantial shared landscape across the isogenic series, with 21,417 loci (70.3%) detected in at least two genotypes, including 11,465 (37.6%) detected in all four, whereas 9,054 loci (29.7%) were detected in only one genotype (Fig. 2B).

Principal component analysis of genome-wide BG4 signal showed clustering of biological replicates and clear separation of the four genotypes. PC1, which explained 47.5% of the variance, primarily separated the BRCA1/2-deficient lines from WT and TP53−/− cells, whereas PC2, explaining 11.2%, further distinguished WT from TP53−/− and TP53−/−;BRCA1−/− from TP53−/−;BRCA2−/− cells (Fig. 2C).

We next examined G4 peak density across genomic annotation classes (Fig. 2D). Relative to the global median, G4 peak density was increased by approximately 780 peaks/Mb at TSSs, 480 peaks/Mb at 5′ UTRs, and 450 peaks/Mb at CpG islands, whereas intronic and intergenic regions were approximately 20 peaks/Mb below the median. After calculating G4 peak density as peaks per megabase of annotated sequence, genotype-specific differences within each annotation class were summarized by median-centering the four genotype values and scaling by the median absolute deviation. TP53−/−;BRCA1−/− cells had the highest relative G4 density in 13 of 14 classes and were tied for the highest in the remaining class, with robust within-region z-scores of 0.7–1.3. In contrast, TP53−/−;BRCA2−/− cells had the lowest relative density across all 14 classes, with robust z-scores ranging from −1.1 down to values at or below −3.0. Given the distinct roles of TP53, BRCA1 and BRCA2 in DNA damage responses,

homologous recombination and replication-stress management, these genotype-associated differences may reflect altered genome-maintenance and replication-stress states in the respective genetic backgrounds.

Functional enrichment analysis of genotype-level G4-associated regions revealed broadly similar functional themes across the four genotypes (Fig. 2E). DNA metabolic process, DNA damage response, and DNA repair were among the strongest enriched terms in each background, with −log10(adjusted P) values of 27.26–32.96, 24.85–30.38, and 21.89–28.06, respectively. Broader enrichment was also observed across the genotypes for chromatin organization and remodeling, transcriptional regulation, RNA processing, and protein catabolic processes. Thus, despite genotype-associated differences in the abundance and genomic distribution of G4 loci, the major functional enrichment themes were similar across the four backgrounds.

At the chromosomal scale, length-normalized G4 peakdensity patterns were largely conserved across the isogenic series (Fig. 2F), although notable differences emerged. Chromosomes 19 and 17 showed densities of 18.39–21.28 and 12.43–17.58 peaks/Mb, respectively, whereas chromosome X showed densities of 1.84–3.19 peaks/Mb. TP53−/−;BRCA2−/− cells had the lowest chromosome-level G4 density among the four genotypes on 21 of 23 chromosomes. Representative loci also showed genotype-associated differences in G4 signal and called peaks (Fig. 2G).

Overall, the chromatin G4 landscape was largely conserved across the isogenic series, with genotype-associated differences in G4 abundance and occupancy superimposed on broadly shared genomic and functional features. These differences may reflect the distinct effects of TP53, BRCA1 and BRCA2 loss on genome maintenance and replication-stress responses, although the present analysis does not establish their underlying mechanisms.

### DNA sequence distinguishes chromatin-associated G4s from loci with in vitro G4-forming potential

We asked whether DNA sequence could distinguish chromatin-associated G4s, identified by consensus BG4 and SG4 CUT&Tag, from sequences with similar composition and intrinsic G4-forming potential. We developed G4SCOPE, a multi-task convolutional–transformer model that jointly predicts G4 occupancy and quantitative CUT&Tag signal from sequence (Fig. 1). The convolutional backbone learns local sequence features that distinguish captured from background regions, while transformer selfattention integrates information across the window and supports positional interpretation of longer-range dependencies. In each genotype, G4-positive loci were evaluated against four background sets of increasing similarity: random genomic windows, GC-matched windows, in vitro only loci defined as G4-seq loci not detected by CUT&Tag in any genotype, and GC-matched in vitro only loci. The last comparison provided the most stringent test because positive and negative loci had similar GC content and evidence of intrinsic G4-forming potential, but differed in whether they were detected as G4s in chromatin. Some in vitro only loci may nevertheless represent CUT&Tag false negatives rather than a complete absence of chromatin folding. Comparisons were performed within each genotype, avoiding direct comparisons of CUT&Tag signal between cell lines (see Methods for background sampling criteria). Models were trained, validated, and tested using non-overlapping chromosome partitions to reduce genomic leakage (Methods). The classifier distinguished G4-positive from background regions in WT, TP53−/−, TP53−/−;BRCA1−/−, and TP53−/−;BRCA2−/− cells (Fig. 3A,B).

**Figure 3.**
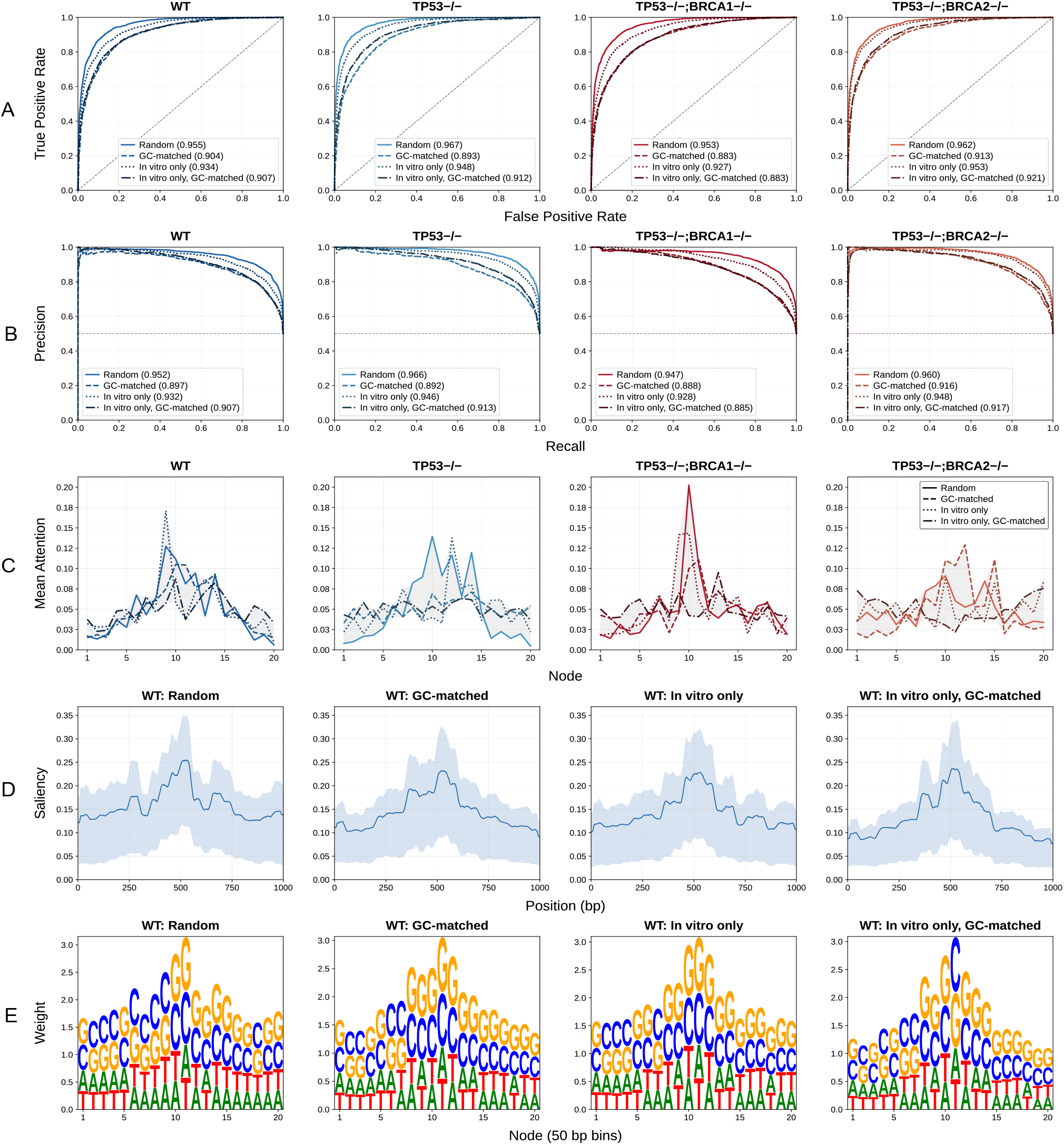
Classification performance and interpretation of G4SCOPE across breast epithelial genotypes. **(A)** Classification performance and interpretation of G4SCOPE across breast epithelial genotypes. Receiver operating characteristic curves for models trained and evaluated in WT, TP53−/−, TP53−/−;BRCA1−/− and TP53−/−;BRCA2−/− cells using four control sets: random, GC-matched, in vitro only (G4-seq loci not detected by CUT&Tag) and in vitro only GC-matched. AUROC values are shown in parentheses. **(B)** Precision–recall curves for the same settings, with AUPRC values shown in parentheses. The horizontal dashed line indicates class prevalence. **(C)** Mean attention weights across the 20 sequence tokens for each genotype and control set, showing increased attention near the centre of the 1-kb input window. Shaded bands indicate the range across control sets. **(D)** Mean saliency profiles for held-out WT sequences, showing predictive importance concentrated near the centred G4 locus. Shaded bands indicate the interquartile range across loci. **(E)** Sequence logos derived from saliency-weighted position frequency matrices for WT sequences, showing enrichment of central G-rich and C-rich patterns.

ROC analysis showed strong discrimination across genotype and background combinations, with AUROC values ranging from 0.889 ± 0.024 to 0.965 ± 0.014 (Fig. 3A; Supplementary Table 1). Performance varied systematically with background definition, with AUROC highest for random negatives, intermediate for in vitro only negatives, and lowest for the more stringent GC-matched and in vitro only GC-matched controls. Performance declined as background loci became increasingly similar to G4-positive loci, but remained strong when chromatin-associated G4s were compared with GC-matched loci possessing in vitro G4-forming potential. This indicates that the model learned sequence features associated with selective G4 occupancy in chromatin rather than relying only on GC content or intrinsic folding potential.

Precision-recall analysis showed the same pattern (Fig. 3B), with AUPRC values ranging from 0.888 ± 0.023 to 0.964 ± 0.014 across genotypes and background definitions (Supplementary Table 1). Given the balanced evaluation sets, AUPRC largely paralleled AUROC. Threshold-dependent performance was similarly stable, with F1 scores ranging from 0.807 ± 0.027 to 0.899 ± 0.018 and accuracies from 0.801 ± 0.028 to 0.897 ± 0.019. Confusion matrices across the 16 genotype-by-background combinations showed consistently more correct than incorrect classifications (Supplementary Fig. 1). Together, these results show that performance was consistent across threshold-free and threshold-dependent measures.

We next assessed whether predicted probabilities corresponded to observed G4-positive frequencies using reliability diagrams and expected calibration error (ECE). Calibration curves remained close to the diagonal across the 16 genotype-by-background combinations, with ECE values ranging from 0.026 to 0.072 (Supplementary Fig. 2). within the balanced evaluation sets used here, predicted probabilities were generally consistent with observed outcome frequencies.

To assess sensitivity to chromosome partitioning, we repeated the classification analysis across three additional predefined whole-chromosome train, validation and test splits while keeping the pipeline unchanged. Performance remained stable across partitions in all four genotypes, with only modest variation in AUROC, AUPRC and F1, indicating that the results were not driven by the primary chromosome split (Supplementary Tables 2 and 3).

Interpretability analyses further showed that predictions were driven mainly by localized sequence features near the centre of the input window, corresponding to the BG4/SG4 CUT&Tag peak (Fig. 3C–E).

Mean attention was concentrated over sequence tokens near the centre, while saliency profiles peaked at the midpoint of the input window, consistent with the position of the G4 locus. Sequence-logo analysis of informative positions also revealed enrichment for G-and C-rich patterns in this region. These results indicate that the classifier learned localized sequence features characteristic of G4-enriched sequences within antibody-captured regions. The motifs identified by the model were highly consistent across negative set background definitions: pairwise cosine similarities between saliency-weighted position frequency matrices ranged from 0.983 to 0.996 (Supplementary Fig. 3). This indicates that the learned sequence features were not driven primarily by background composition.

### Sequence-based modeling recapitulates quantitative variation in G4 signal intensity

Having shown that the classifier component of the multitask model could identify CUT&Tag-defined G4-associated regions from DNA sequence, we next asked whether the same jointly trained model could also capture quantitative CUT&Tag signal profiles, in addition to distinguishing antibody-captured regions from background regions. Signal reconstruction tests whether the shared sequence representation also captures the shape and magnitude of local enrichment. The classification and signal-reconstruction heads were optimized simultaneously within a single multi-task training procedure, allowing us to evaluate whether the shared representation supports both tasks. In randomly chosen held-out windows, predicted profiles recapitulated the measured signal distribution across antibody-captured regions in all four genotypes, capturing the position, extent, and relative magnitude of enrichment (Fig. 4A). In contrast, predicted signal remained low across background regions, indicating that the model learned not only to identify enriched loci but also to distinguish them from windows with weak or flat signal profiles. Although predicted profiles were generally smoother than the measured signal, they preserved the locus-level trends in signal intensity and shape.

**Figure 4.**
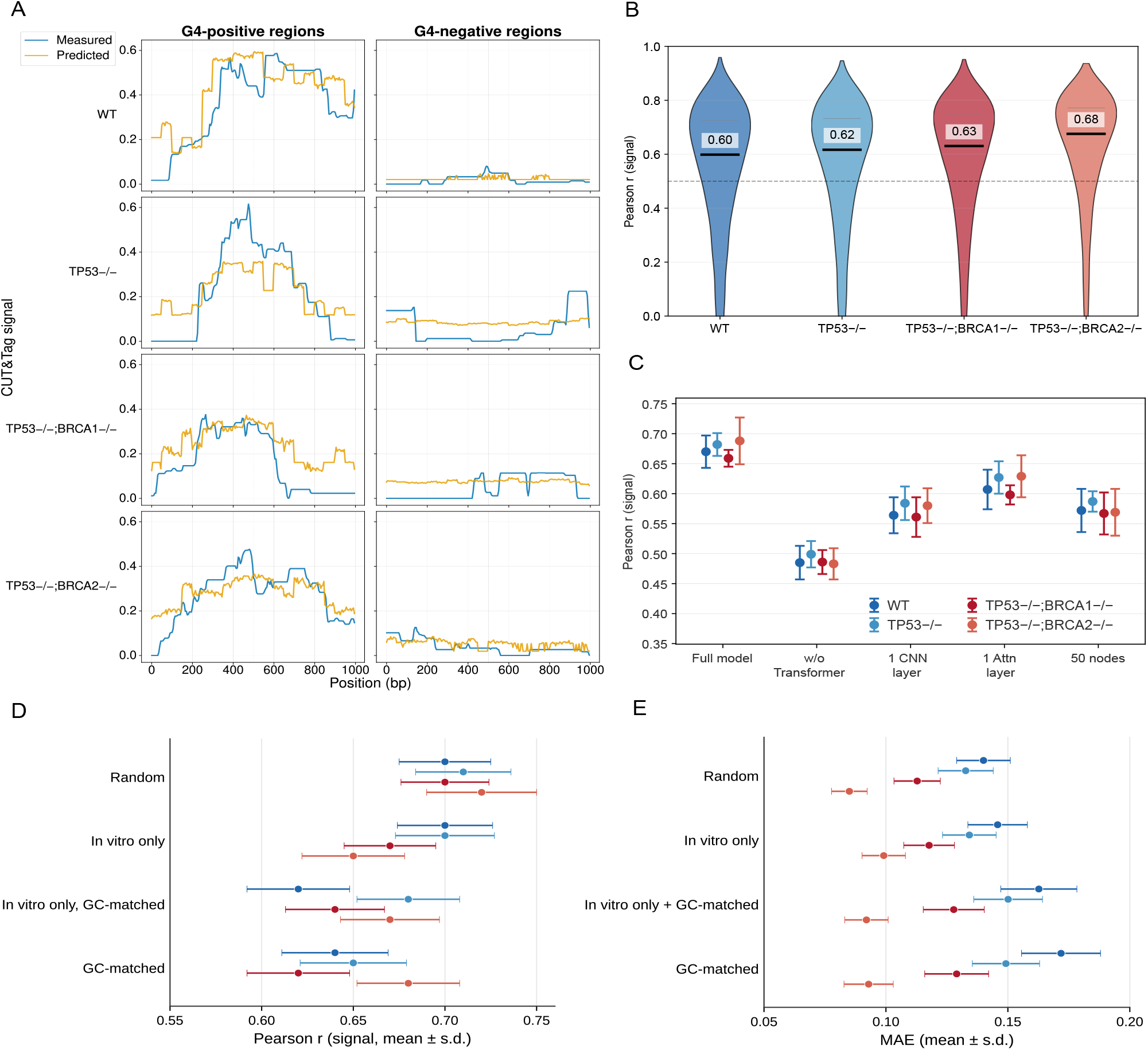
Quantitative CUT&Tag signal prediction and architectural ablation across breast epithelial genotypes. **(A)** Representative measured (blue) and predicted (orange) CUT&Tag signal profiles for G4-positive (left) and G4-negative (right) genomic windows in WT, TP53−/−, TP53−/−;BRCA1−/− and TP53−/−;BRCA2−/− cells. The vertical dashed line marks the centre of the input window. **(B)** Distribution of per-window Pearson correlation coefficients between predicted and observed signal profiles for held-out G4-positive windows for which Pearson correlation was defined, pooled across background definitions. Bold horizontal lines and numeric labels indicate medians, thin grey lines indicate the 75th percentile and the dashed line marks r = 0.5. **(C)** Ablation analysis of signal-prediction performance. Mean Pearson correlation coefficients are shown for the full model and four architectural variants: removal of the Transformer encoder, reduction to a single CNN layer, reduction to a single attention layer and use of a 50-node sequence representation. **(D)** Mean Pearson correlation coefficients for each genotype using four negativeset definitions: random, in vitro only, in vitro only GC-matched and GC-matched. **(E)** Mean absolute error (MAE) for the same genotype and negative-set combinations shown in **(D).** Colours in **(C-E)** indicate genotype. Error bars indicate ±1 s.d. across three random seeds.

Similar agreement was observed across the held-out set. Among held-out G4-positive windows for which Pearson correlation between the predicted and observed signal profiles was defined, median correlations were 0.60 for WT, 0.62 for TP53−/−, 0.63 for TP53−/−;BRCA1−/−, and 0.68 for TP53−/−;BRCA2−/− (Fig. 4B).

Genotype- and background-set-level summary metrics showed the same overall pattern. Mean Pearson correlation ranged from 0.62 ± 0.028 to 0.72 ± 0.030, mean absolute error from 0.085 ± 0.007 to 0.172 ± 0.016, and mean squared error from 0.026 ± 0.003 to 0.079 ± 0.009 (Fig. 4D,E; Supplementary Table 4). As in the classification task, performance generally decreased with increasing background stringency, with the highest correlations typically observed for random negatives and lower performance for GC-matched or in vitro only controls. Despite this increase in difficulty, regression performance remained robust across all genotypes and background definitions.

Performance was broadly consistent across the isogenic series, with the strongest overall agreement observed in TP53−/−;BRCA2−/−, which showed the highest median per-locus correlation in the held-out set, the highest or near-highest mean Pearson correlations across negative-set controls, and the lowest MAE overall (Fig. 4B,D,E; Supplementary Table 4). Together, these results show that the multi-task architecture supports sequencebased prediction of G4 presence and fine-scale variation in CUT&Tag signal intensity.

We next examined cross-genotype transfer by evaluating each genotype-trained model on held-out chromosomes from the other genotypes. Performance remained strong in both matched and cross-genotype settings, with only modest differences between within-genotype and cross-genotype evaluations (Supplementary Fig. 4). This suggests that the major sequence features associated with chromatin G4 occupancy are shared across the mammary epithelial genotypes. Whether an already-trained mammary epithelial model transfers across cellular lineages remains unresolved; however, we assessed the portability of the G4SCOPE framework by independently applying it to a published K562 BG4 CUT&Tag dataset.

### Ablation studies define the contributions of convolutional feature extraction, contextual self-attention and multi-task learning

To define the contribution of individual architectural components, we performed ablation studies in which components of the full model were systematically removed or modified and compared across genotypes and negative-set definitions (Fig. 4C; Supplementary Fig. 5; Supplementary Tables 5–8; see Methods). The full model achieved the highest AUROC and signal correlation across all settings. Removing the Transformer caused the largest decrease in performance, reducing AUROC by 12.1 to 13.2% and signal correlation by 26.3 to 29.8%. Simplifying the CNN backbone to a single convolutional layer also impaired performance, reducing AUROC by 9.5 to 9.6% and signal correlation by 12.9 to 14.3%. Removing the auxiliary signal-prediction task reduced classification performance by 7.2 to 7.5%, indicating that the auxiliary signal-prediction task improves occupancy classification by providing complementary supervision over the shared G4-associated sequence representation. Additional architectural variants, including reducing the model to a single attention layer or increasing the number of sequence tokens also yielded lower classification and signal-prediction performance than the full model.

**Figure 5.**
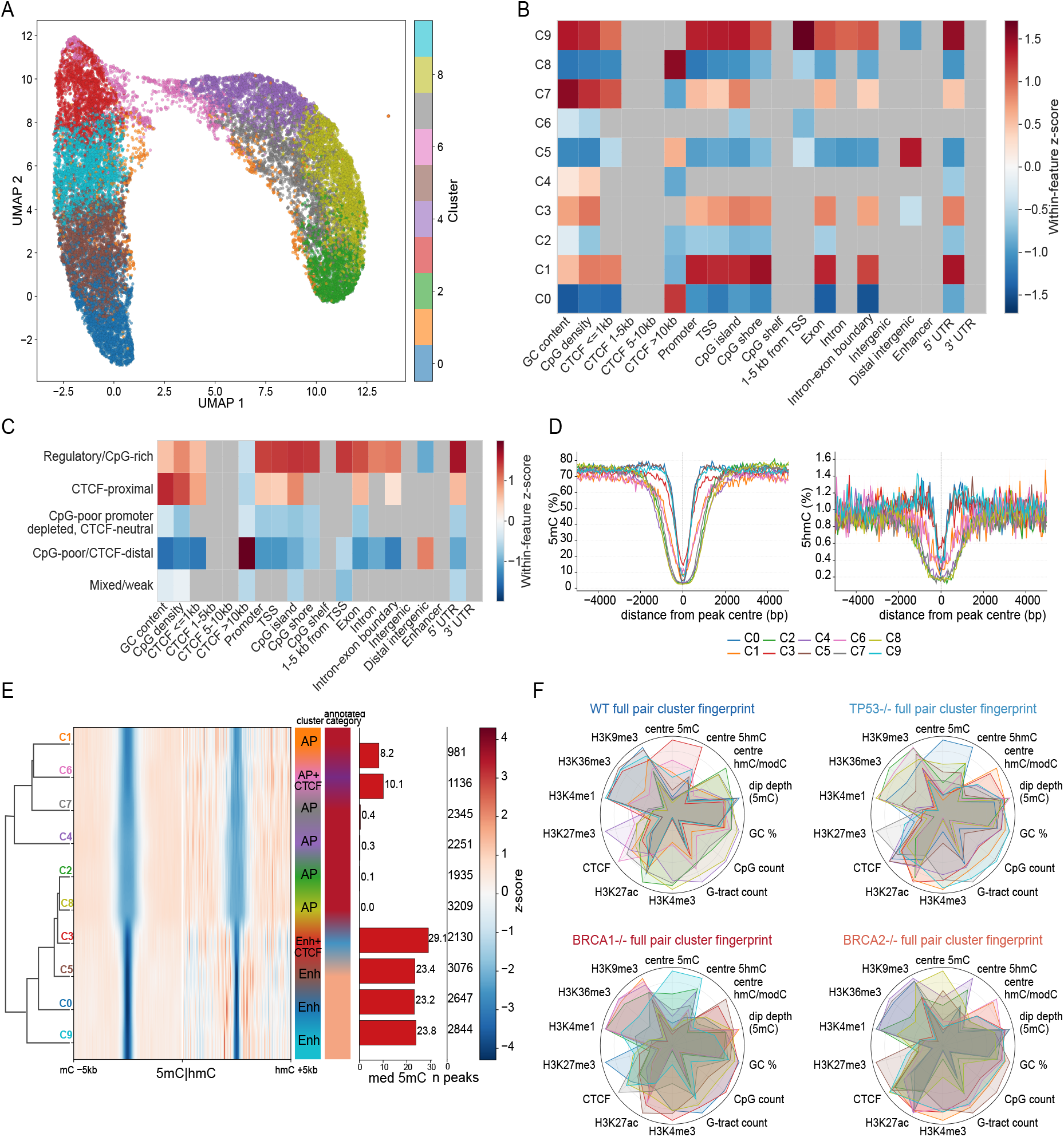
Sequence-defined G4 subclasses occupy distinct regulatory and cytosine-modification environments. **(A)** UMAP visualization of model-derived G4 embeddings coloured by cluster assignment. **(B)** Within-feature standardized enrichment of sequence and genomic annotations across WT G4 embedding clusters. Categorical annotations were evaluated using Fisher’s exact test and continuous features using the Mann–Whitney test, with Benjamini–Hochberg correction for multiple testing; grey indicates associations with FDR ≥ 0.05. **(C)** Enrichment profiles after grouping clusters with similar annotation patterns into five biologically interpretable meta-clusters. **(D)** Cluster-specific 5mC and 5hmC profiles centred on G4 loci. **(E)** Hierarchical clustering and row-standardized heatmaps of 5mC and 5hmC profiles for each cluster, shown together with the dominant cCRE class, median 5mC and number of peaks. **(F)** Radar plots summarizing sequence, cytosine-modification and chromatin-feature fingerprints for WT, TP53−/−, TP53−/−;BRCA1−/− and TP53−/−;BRCA2−/− G4 cluster sets.

### Benchmarking and independent evaluation support modeling chromatin-associated G4s using cell line data

We next benchmarked our classifier against established G4 prediction methods using the stringent in vitro only GC-matched benchmark. To ensure a matched comparison, trainable baseline classifiers were trained from scratch on the same genotype-specific CUT&Tag datasets, and evaluated using the same chromosome-holdout framework as our model (see Methods). Implementation details for the benchmarked methods are provided in Supplementary Table 9. In this setting, G4SCOPE matched or exceeded the established classifiers, with mean AUROC, AUPRC and F1 values of 0.912 ± 0.012, 0.913 ± 0.010 and 0.837 ± 0.012, respectively (Supplementary Table 10). DeepG4 and G4-Attention were the strongest comparison methods, with G4-Attention achieving a similar mean F1 of 0.827 ± 0.013. Unlike these classifiers, G4SCOPE was jointly trained to predict occupancy and quantitative CUT&Tag signal. By contrast, the rule-based methods pqsfinder, Quadparser, and G4Hunter performed near chance, indicating that motif-based scoring had limited discriminatory power when control examples were restricted to in vitro only GC-matched loci. This suggests that discrimination of chromatin-folded G4 loci depends on sequence features beyond core motif composition, potentially reflecting flanking sequence context associated with chromatin accessibility or folding propensity.

At the genotype level, our model remained consistently strong across WT, TP53−/−, TP53−/−;BRCA1−/− and TP53−/−;BRCA2−/−, with AUROC values ranging from 0.900 ± 0.023 to 0.927 ± 0.020 and AUPRC values ranging from 0.903 ± 0.022 to 0.926 ± 0.020 (Supplementary Tables 11 and 12). Performance was highest in TP53−/−;BRCA2−/−, a trend observed for several retrained baselines. Genotype-level F1 values showed the same overall pattern, with our model achieving the highest thresholded performance across all genotypes and G4-Attention remaining the closest baseline (Supplementary Table 13). Model complexity and inference-time comparisons are provided in Supplementary Table 14.

G4mismatch performed poorly across all genotypes, with mean AUROC and AUPRC values of 0.202 ± 0.052 and 0.353 ± 0.030, respectively, and genotype-specific AUROCs ranging from 0.170 to 0.276 (Supplementary Tables 10–12). Because G4mismatch was included as a zero-shot transfer baseline trained on G4-seq-derived labels, and was not calibrated using CUT&Tag-specific thresholds, we report only threshold-free metrics. Its performance suggests that this G4-seq-trained model did not transfer to CUT&Tag-defined G4 occupancy in the 184-hTERT system, although this may reflect assay differences or cell-context-specific sequence grammar. Quantitative signal regression was not benchmarked against the comparison methods, as none of the baselines were designed to predict CUT&Tag signal intensity in this setting. Overall, the model outperformed the retrained classifiers, rule-based methods and zero-shot transfer baseline on genotype-specific chromatin G4 occupancy.

To determine whether the G4SCOPE framework could be applied beyond the 184-hTERT breast epithelial system, we independently trained and evaluated the same architecture using a published K562 BG4 CUT&Tag dataset (GSE181373) and the same chromosome-holdout framework. This analysis tested the portability of the modeling framework to another cellular lineage rather than zero-shot transfer of a model trained in breast epithelial cells. On the stringent in vitro only GC-matched benchmark, the model achieved an AUROC of 0.918, an AUPRC of 0.907 and an F1 score of 0.841, while outperforming the learned and rule-based baselines (Supplementary Tables 15 and 16). It also predicted quantitative CUT&Tag signal on held-out chromosomes with a Pearson correlation of 0.618. These results show that independent application of the G4SCOPE framework supports both occupancy and quantitative signal prediction in a distinct cellular lineage.

### Sequence-defined G4 subclasses occupy distinct regulatory and cytosine-modification environments

Having established that sequence distinguishes chromatin-associated G4s from loci with intrinsic G4-forming potential, we asked whether the learned sequence representations also delineate biologically distinct classes of chromatin G4s. We therefore reduced attention-weighted G4SCOPE embeddings from BG4/SG4-captured loci using PCA and clustered the resulting representations using Gaussian mixture modelling. BIC-based model selection identified 10 clusters (Supplementary Fig. 6), which were represented across all cell lines. UMAP was used independently to visualize the resulting cluster structure in two dimensions (Fig. 5A; Methods), motivating further characterization of their sequence, genomic and epigenetic features.

**Figure 6.**
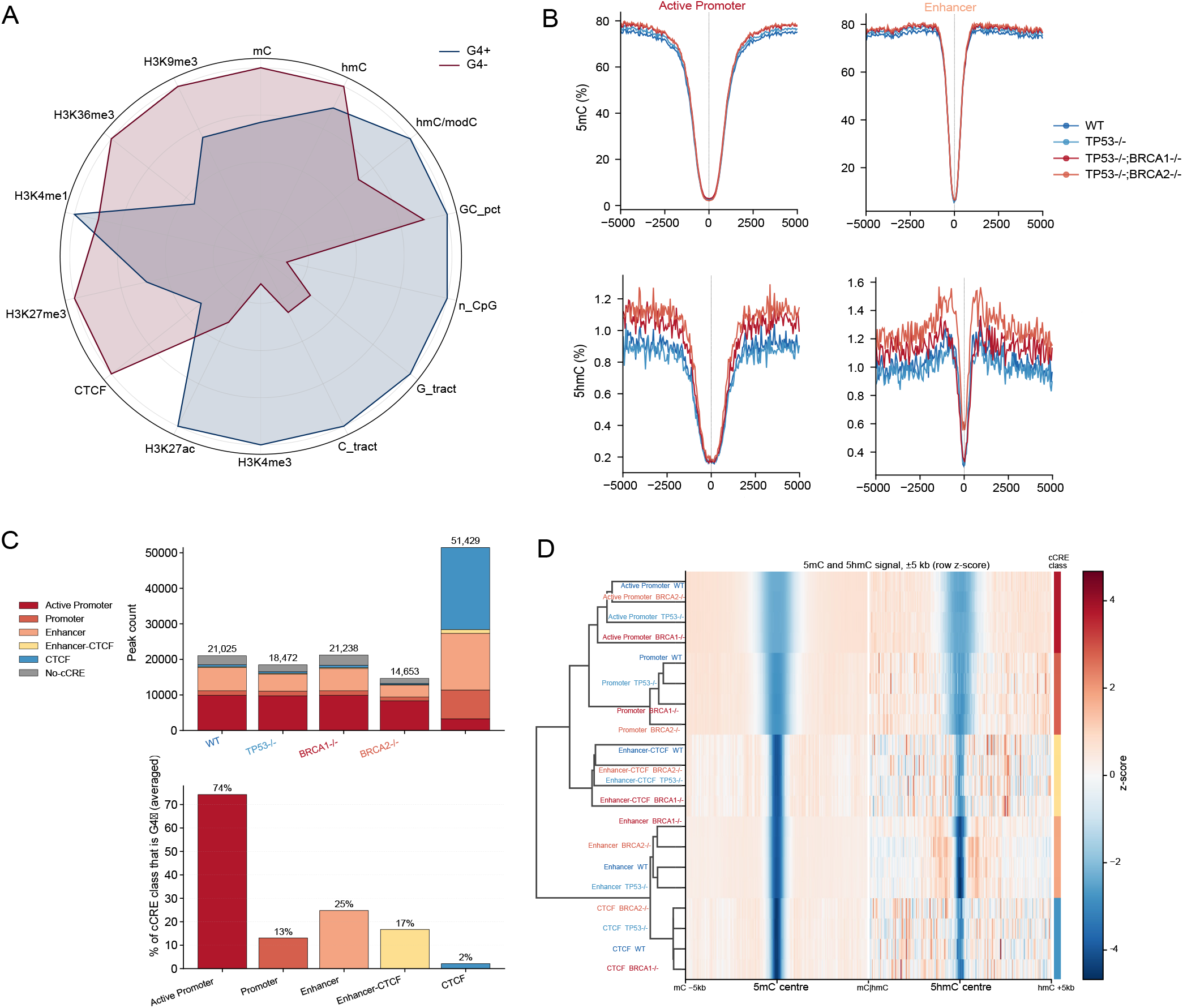
Chromatin-associated G4s delineate regulatory and cytosine-modification environments. **(A)** Radar plot comparing sequence, cytosine-modification and chromatin features between experimentally defined G4-positive (G4+) and G4-negative (G4−) regions, including GC and CpG content, G-and C-tracts, 5mC, 5hmC, histone marks and CTCF. **(B)** 5mC (top) and 5hmC (bottom) metaplot profiles centred on G4 loci within active-promoter and enhancer cCREs across the four genotypes. **(C)** Distribution of G4-positive peaks across cCRE classes in each genotype and in the union set (top), and the percentage of sites within each cCRE class overlapped by G4 peaks, averaged across cell lines (bottom).**(D)** Hierarchical clustering of row-z-scored 5mC and 5hmC profiles within ±5 kb of G4 centres across genotypes and cCRE classes, together with the corresponding cCRE class annotation.

We first examined whether the embedding-derived clusters were associated with distinct sequence and genomic contexts, including GC and CpG content, regulatory and gene-structural annotations, and proximity to CTCF-binding sites. In WT cells, the clusters showed marked differences across these features (Fig. 5B). Several clusters were CpG-and GC-rich and enriched at CpG islands, promoters and TSS-associated regions, whereas others were CpG-poor and depleted from promoter-associated regions. CTCF proximity also differed among clusters, with one cluster enriched for CTCF-proximal loci and several CpG-poor clusters associated with CTCF-distal contexts. Grouping clusters with similar enrichment patterns yielded five broader meta-clusters representing Regulatory/CpG-rich, CTCF-proximal, CpG-poor promoter-depleted/CTCF-neutral, CpG-poor/CTCF-distal and Mixed/weak contexts (Fig. 5C). We then examined whether these clusters showed distinct transcription-factor motif patterns. Although GC-box and KLF-family motifs were enriched across clusters, their positional distributions differed, with the strongest cluster-specific differences observed for AP-1-family motifs (Supplementary Fig. 7). More broadly, each cluster showed a distinct motif signature, suggesting differences in regulatory context (Supplementary Fig. 8). Given prior evidence linking G4 structures to local DNA methylation landscapes ^28^, we next asked whether the sequence-derived clusters were also associated with distinct cytosine-modification states. To measure these modifications directly in the same isogenic breast epithelial cells, we profiled 5mC and 5hmC by 6-base epigenetic whole-genome sequencing^49^. Cluster-specific 5mC and 5hmC profiles showed distinct patterns around G4 centers in WT cells (Fig. 5D), namely different minimum and distribution of the modifications around the centre. These differences were further resolved by hierarchical clustering, which linked distinct cytosine-modification profiles with differences in regulatory annotation and median 5mC levels across clusters (Fig. 5E). Across genotypes, the resulting sequence and epigenetic feature fingerprints were broadly similar, including comparable CpG-and GC-associated characteristics, with only modest differences between cell lines (Fig. 5F). Together, these results identify sequence-defined subclasses of chromatin G4s that occupy distinct regulatory environments and are associated with characteristic local 5mC and 5hmC profiles.

### Chromatin-associated G4s delineate conserved regulatory and cytosine-modification environments across genotypes

Having identified sequence-defined G4 subclasses with distinct local cytosine-modification profiles, we next asked whether experimentally captured G4 loci occupy conserved regulatory and epigenetic environments across the isogenic series. Using the same 6-base epigenetic whole-genome sequencing data together with publicly available human mammary epithelial cells (HMEC) cis-regulatory elements and histone profiles (ENCSR725DZR), we characterized the regulatory and cytosine-modification environments surrounding G4-positive loci.

In WT cells, G4-positive and G4-negative regions showed distinct sequence, cytosine-modification and chromatin-feature profiles (Fig. 6A). Relative to G4-negative regions, G4-positive loci showed higher H3K4me3 and H3K27ac and lower H3K36me3, CTCF and 5mC-associated feature values, consistent with their association with active regulatory chromatin^26^. By contrast, H3K4me1- and 5hmC-associated regions showed more heterogeneous patterns of G4 occupancy (Fig. 6A). Across the isogenic series, the fraction of sites overlapped by G4 peaks differed across HMEC candidate cis-regulatory element (cCRE) classes and was highest for active-promoter cCREs (Fig. 6C). G4-positive loci were also generally locally hypomethylated, with sharp decreases in both 5mC and 5hmC around G4 centers. When examined within regulatory contexts, the shape of these cytosine-modification profiles differed between active promoters and enhancers, with broader 5mC and 5hmC profiles around active promoters than around enhancers (Fig. 6B). Hierarchical comparison of these profiles further showed that regulatory-element-associated 5mC and 5hmC patterns were broadly conserved across the four genotypes (Fig. 6D). Together, these findings establish chromatin-associated G4s as markers of active regulatory environments with characteristic local 5mC and 5hmC profiles, relationships that are broadly conserved across the isogenic series.

## Discussion

A key goal for epigenomic studies and the detection of secondary-structure-forming regions in the genome is the development of quantitative models that can predict these features from sequence alone. Here, we developed a multitask sequence model that jointly predicts chromatin-associated G4 occupancy and quantitative CUT&Tag signal intensity from a shared sequence representation. Whereas many previous G4 prediction models are binary classifiers or require auxiliary epigenomic inputs such as ATAC-seq, G4SCOPE predicts both occupancy and quantitative signal profiles from DNA sequence alone. Convolutional layers capture local motif-level features, while token-level selfattention integrates sequence context across the full input window. Both G4 occupancy and local signal profiles could be predicted from DNA sequence across an isogenic breast epithelial system comprising WT, TP53−/−, TP53−/−;BRCA1−/− and TP53−/−;BRCA2−/− genotypes.

Performance remained strong when chromatin-associated G4s were compared with GC-matched loci that had in vitro G4-forming potential, indicating that the model learned sequence features beyond broad GC composition or intrinsic folding potential. Cross-genotype performance was also strong, suggesting that much of the sequence information associated with chromatin G4 occupancy is shared across the isogenic series, although this transferability likely reflects, in part, their common mammary epithelial background. Independent training and evaluation in K562 cells showed that the same framework could also be applied in a distinct cellular context. Together, these results support training directly on cell-based G4 data rather than relying solely on canonical sequence motifs or models trained on in vitro G4-seq data.

Our development of G4SCOPE demonstrates that a shared sequence-based encoder can successfully decouple endogenous, chromatin-constrained G4 occupancy from intrinsic, in vitro folding potential^37,39^. While traditional rule-based models struggle when confronted with GC-matched in vitro controls^29,30^, G4SCOPE identifies localized, fine-scale sequence elements that govern cellular folding dynamics. Crucially, by executing quantitative signal profile estimation as a parallel multi-task learning objective, the framework captures structural features that correlate tightly with antibody signal intensity and regulatory context rather than broad sequence composition alone.

Although the overall G4 landscape was largely shared across the isogenic series, we also observed clear genotype-associated differences. TP53−/−;BRCA1−/− cells had the highest relative G4 density across most genomic annotations, whereas TP53−/−;BRCA2−/− cells had the lowest. A particularly striking biological finding from our genotype-resolved mapping is the profound divergence in G4 landscapes between BRCA1- and BRCA2-deficient cells. While TP53−/−;BRCA1−/− contexts exhibit an elevated global density of G4 signals, TP53−/−;BRCA2−/− cells exhibit a widespread, systemic depletion of chromatin-associated G4 structures across nearly all genomic tracks. This phenomenon suggests that although both proteins are vital components of homologous recombination and fork stabilization pathways^19^, the absolute loss of BRCA2 imposes distinct selective pressures on the persistence or clearance of folded secondary structures during replication stress^17^. The genomes of the BRCA1 and BRCA2 isogenic knockouts used here have been previously shown to exhibit the characteristic mutational patterns of HR deficiency as well as BRCA1- and BRCA2-specific mutational patterns^45^.

These global differences may influence the duplication and deletion of regions containing G4-forming sites. The regression results indicate that sequence features are associated not only with whether a G4 is detected, but also with the position, shape and relative intensity of the CUT&Tag signal. However, G4 CUT&Tag signal is likely influenced by several additional factors, including local chromatin and assay accessibility, regulatory context, copy number and genotype-specific cellular state. Therefore, although the model receives sequence alone as input, the signal profiles it learns may reflect sequence features that interact with these broader epigenetic and genomic contexts.

Clustering the model-derived G4 representations identified groups of loci with distinct genomic and epigenetic properties. In the 184-hTERT system, these clusters differed in GC and CpG content, regulatory annotations and CTCF proximity, and were associated with distinct experimentally measured 5mC and 5hmC profiles. Notably, G4SCOPE did not receive 5mC or 5hmC data as input, indicating that these associations emerged from sequence-derived representations learned by the model. The spatial extent of these cytosine-modification profiles also differed by regulatory context, with broader 5mC and 5hmC profiles around G4s at active promoters and narrower, more localized profiles around enhancer-associated G4s. This distinction may reflect differences in CpG density and chromatin organization between promoter and enhancer regions. Transcription-factor motif analysis further revealed cluster-specific motif patterns, linking the sequence features learned by the model to differences in regulatory and cytosine-modification context. Our findings extend previous work by resolving these relationships across sequence-defined G4 subclasses in the same isogenic system.

Furthermore, G4SCOPE’s learned embeddings reveal that chromatin-embedded G4s partition into highly structured sub-populations exhibiting unique transcription factor motif signatures and distinct cytosine modification dynamics. The sharp, localized depletions of 5mC and 5hmC observed around specific G4 centers validate and extend prior models wherein G4 arrays serve as physical barriers to DNA methyltransferases^28^. By demonstrating that these 5mC/5hmC profiles vary symmetrically and topologically across sequence-defined subclasses, our data suggest a highly coordinated interplay where primary sequence grammar simultaneously governs G4 structural topology and local epigenetic remodeling. Further work will be needed to understand these relationships, particularly the association between G4s and 5hmC.

Overall, G4SCOPE provides a framework for predicting chromatin-associated G4 occupancy and quantitative signal from sequence. Its learned representations also provide a basis for examining how G4 sequence features interact with epigenetic, genomic and cell-state factors in other cancer-relevant systems. We believe this architecture will be extensible to multiple modalities and applicable to other CUT&Tag-based profiling methods for assessing epigenetic structure and drug binding.

## Acknowledgements

We thank the members of the Aparicio and Balasubramanian laboratories for insightful discussions. We thank Tehmina Masud for generating the mutant cell lines used in this study. We also thank the staff of the Genomics Core, Research Instrumentation, and Cell Services core facilities at the Cancer Research UK Cambridge Institute for their support. This project was supported by the BC Cancer Foundation at BC Cancer and by the Office of the Assistant Secretary of Defense for Health Affairs through the Breast Cancer Research Program under Award No. HT9425-23-1-0820. S.A. holds the Nan and Lorraine Robertson Chair in Breast Cancer and is a Canada Research Chair in Molecular Oncology (950-230610 and CRC-2021-00205). This work was also supported by the Canadian Cancer Society Research Institute Impact Program grant 705617, Canadian Institutes of Health Research grant PJT-190058, Breast Cancer Research Foundation award BCRF-24-180, Cancer Research UK core funding (C9545/A29580; DRCQQRNov21/100001), Cancer Research UK programme funding (C9545/A19836), and Herchel Smith funds from the University of Cambridge.

## Author contributions

Conceptualization: A.S., I.E.-G., L.M., S.B., S.A.; Methodology: A.S., I.E.-G., L.M.; Software: A.S.; Formal analysis: A.S., I.E.-G., L.M.; Investigation: A.S., I.E.-G., L.M., D.Y., C.O., V.C., D.L.; Data curation: A.S., I.E.-G., L.M.; Validation: A.S.; Visualization: A.S., I.E.-G., L.M.; Funding acquisition: S.B., S.A.; Supervision: D.T., Y.P., S.B., S.A.; Writing – original draft: A.S., I.E.-G., L.M.; Writing – review & editing: A.S., I.E.-G., L.M., S.B., S.A., with contributions from all authors..

## Data and materials availability

The sequencing data generated in this study have been deposited in the European Genome-phenome Archive (EGA) under study accession EGAS50000002094. G4 CUT&Tag sequencing data are available under dataset accession EGAD50000003011, and biomodal 6-base sequencing data are available under dataset accession EGAD50000003012. Access to these datasets is controlled by the BC Cancer/Provincial Health Services Authority Data Access Committee and requires an executed Data Access Agreement; access requests can be initiated through the EGA dataset pages. The published K562 BG4 CUT&Tag data used in this study are publicly available from the Gene Expression Omnibus under accession GSE181373. Cisregulatory elements and histone modification profiles were annotated using data from the ENCODE project ENCSR725DZR. Code for training and evaluating the multi-task G4 prediction model, including data preprocessing, model architecture, ablation analyses, and benchmarking, is available at https://github.com/molonc/multi-task-G4-prediction.

## Materials and methods

### Human mammary epithelial cell culture

The wild-type hTERT-immortalized non-transformed human mammary epithelial cell line 184-hTERT L9^44^ and isogenic derivatives (TP53−/−, TP53−/−;BRCA1−/− and TP53−/−;BRCA2−/−) were cultured as previously described in mammary epithelial basal medium (MEBM; Lonza) supplemented with the SingleQuots additives (Lonza), 5 µg ml−1 transferrin (Sigma-Aldrich) and 10 µM isoproterenol (Sigma-Aldrich)^45^. Cell lines were routinely confirmed mycoplasma-free and authenticated by STR profiling and allele genotyping. Cells of near-equivalent passages were selected for experimentation.

### G4 mapping using CUT&Tag

BG4 (a recombinant, FLAG-tagged single-chain variable fragment, scFv, antibody that recognizes folded DNA G4 structures) and SG4 (a recombinant single-domain nanobody targeting G4s) were expressed, purified, and validated as previously described^13^. In situ mapping was performed as previously described^47,48^ on 100,000 cells. Cells were fixed and immobilized on ConA-conjugated magnetic beads. BG4 was added (1:50) followed by incubation at 4 °C overnight at 600 rpm. Anti-flag secondary antibody was added (1:50) and samples incubated at 25 °C for 2 h at 600 rpm. Tertiary antibody (1:100) was added followed by pA-Tn5 (1:250) and samples incubated at 25 °C for 1 h at 600 rpm. Tagmented DNA was purified with DNA clean & concentrator (Zymo Research, cat #D4013). Sequencing libraries were generated as previously described using universal i5 and uniquely barcoded i7 primers^13,50^. Sequencing was performed on an Illumina NextSeq 2000 using either a High Output Kit v2.5 with 75 cycles or P2/P3 reagents with 100 cycles. No-primary antibody control libraries did not yield sufficient material for sequencing.

### 6-base (5mC and 5hmC) whole-genome library preparation and data processing

The duet evoC assay (biomodal) was carried out according to the manufacturer’s recommendations. A control DNA spike-in was included to evaluate 5mC/5hmC conversion and library preparation efficiency. Genomic DNA was extracted using the DNeasy Blood and Tissue kit (Qiagen) and fragmented to ∼250bp using the Covaris M220 ultrasonicator. 80ng of sheared DNA was end-repaired, A-tailed, and ligated to hairpin adapters. Hairpins were then digested, a complementary strand was copied, followed by forkhead adapter ligation and copying of 5mCs. 5hmCs were then oxidized, followed by deamination of unmodified cytosines. The libraries were then PCR amplified to include unique indices for Illumina sequencing. Final libraries were analyzed by Qubit and Agilent Bioanalyzer and pooled at equimolar concentration. Illumina sequencing was performed on a NovaSeqX Plus at the Genome Sciences Centre, Vancouver, BC to generate 2 x 150bp reads, with 5-15% PhiX spike-in. Median sequencing depth was 30x. 6-base sequencing data were processed with the biomodal duet software (v1.3.0) in Nextflow (v24.04.3), and reads were aligned with BWA-MEM2.

### CUT&Tag sequencing data processing

Illumina sequencing paired-end output files were demultiplexed using demux Illumina version 3.0.9 using the flags; -c -d -i -e -t 1 -r 0.01 -R -l 9. The resulting fq.gz files underwent sequencing quality control using FastQC v0.11.8, and their summary was visualized by MultiQC v1.11. Bases with a quality score below 20 were trimmed from both reads using cutadapt (cutadapt -q 20). Fastq files were aligned to the combined hg38 and *E. coli* genomes (or the corresponding genome builds which include the modified site confirmed by Sanger sequencing) using bwa 0.7.17-r1188 with only reads in the whitelist regions of hg38 continuing the process pipeline. Duplicates were removed using Picard version 2.20.3 (Picard MarkDuplicates). Peaks were called using SEACR version 1.3, without input control, reporting the top 1% by area under curve (AUC) regions, using both the relaxed and stringent criteria. Unless otherwise stated, downstream analyses used SEACR stringent peak calls. For each genotype and probe, replicate-consensus peaks were defined as peaks detected in at least two of three biological replicates (≥2/3). Assessment of peak overlap across biological replicates was performed using intervene upset and pairwise modules. High reproducibility across samples was confirmed. BigWig files were created using deepTools version 2.0 bamCoverage using the mapped BAM files and the flags --binSize 3 --normalizeUsing CPM -- extendReads, for fragments of up to 600 nt (using –maxFragmentLength).

### Generation of hg38 genomic annotation tracks

Genomic annotation tracks were assembled from public reference resources and standardized to BED (0-based, half-open) coordinates. Gene features were parsed from the Ensembl GRCh38 GTF (release 112) and restricted to canonical autosomes and chrX ^51^. Exons and gene bodies were merged into interval sets, with introns defined as gene bodies minus exons. Strand-specific coordinates were used to derive TSS (±500 bp), promoters (−2 kb/+500 bp), and 1–5 kb upstream regions; exon boundaries were defined as ±200 bp around exon starts and ends. UTRs were extracted from GTF annotations and classified as 5′ or 3′ relative to CDS boundaries. CpG islands were obtained from UCSC (cpgIslandExt) and extended to shores (0–2 kb) and shelves (2–4 kb) ^52^. Enhancers were defined using ENCODE cCREs with enhancer-like signatures (pELS/dELS) ^53^. All tracks were sorted and merged to generate non-redundant BED files for overlap-based annotation.

### Chromosomal and genomic annotation distributions of G4 peaks

To quantify patterns of shared and genotype-restricted G4 detection, we performed an overlap analysis using the replicate-consensus peak sets defined above. Peak intervals from all genotypes were merged to define a unified set of genomic loci by collapsing overlapping intervals into single representative peaks. For each merged locus, presence in each genotype was assessed against the original peak calls, and a peak was considered present when overlap was at least 500 bp, to capture positional concordance while excluding marginal edge overlaps.

To summarize G4 occupancy at the chromosome scale, peaks were counted per chromosome (chr1–chr22 and chrX) for each genotype and normalized by chromosome length to obtain peak density (peaks/Mb). For relative comparison across chromosomes and genotypes, densities were centered by subtracting the global median across all finite chromosome-by-genotype values.

To characterize G4 distribution across genomic annotation classes, G4 density was calculated as peaks per megabase of annotated sequence for each region category and genotype. Relative genotype-specific differences within each annotation class were summarized using robust row-wise standardization, in which values were centered by the median and scaled by the median absolute deviation. Overall region-level enrichment or depletion was assessed by comparing the per-region central tendency across genotypes with the global median of the full region-by-genotype density matrix.

### Training dataset construction and background selection

SEACR peak calling and replicate-consensus peak definition were performed genome-wide before model dataset construction and independently of the chromosome-based train, validation and test partitioning. For deep learning-based G4 prediction, genotype-specific datasets were constructed from the human reference genome at a 1:1 ratio of positive to background windows. This design enabled controlled comparisons rather than reproducing genome-wide G4 prevalence. Analyses were performed separately for each of the four isogenic genotypes. All examples were represented as fixed 1 kb windows. For each genotype, positive examples were derived from its corresponding CUT&Tag peak calls by centering a 1 kb window on each peak midpoint (±500 bp). The merged peak set was used only for overlap analyses and not for model training. Windows extending beyond chromosome boundaries or containing ambiguous bases (N) in hg38 were excluded. For each retained window, the DNA sequence was extracted and one-hot encoded as a 1,000 × 4 matrix representing A, C, G and T.

For each genotype, background examples were generated using four complementary sampling strategies to evaluate robustness to negative-set definition. Background coordinates were sampled separately for each genotype, and their quantitative signal targets were derived from the corresponding genotype-specific CUT&Tag track. All negative windows were required to be 1 kb, chromosome-boundary safe, free of ambiguous bases, non-overlapping with one another within a given negative set, and to have zero overlap with the union of positive windows across all genotypes. Random negatives were sampled genome-wide using per-chromosome quotas matched to the positive count distribution for each genotype. GC-matched negatives were sampled from the genome to match the GC-content distribution of the corresponding positive set. In vitro only negatives were derived from published G4-seq loci by generating 1 kb windows centered on each site and retaining only loci that did not overlap any CUT&Tag-positive window in any genotype. A fourth set, in vitro only GC-matched, comprised GC-matched negatives generated from these in vitro only candidates.

For GC-matched sampling, GC content was calculated for each 1 kb window as (G + C)/1000 after excluding windows containing N. Negatives were generated at 1:1 parity with positives and matched within chromosome using an expanding GC tolerance of ±0.005, ±0.010, ±0.020, ±0.030, ±0.050 and ±0.080 relative to each positive window. When no admissible match was identified within these tolerances, a nearest-GC fallback was applied by sampling a large pool of eligible windows from the same chromosome and selecting the window with the smallest absolute GC difference. Sampling used a fixed random seed for reproducibility.

### Multi-task model architecture, training and evaluation

We developed G4SCOPE, a multi-task deep learning framework that predicts both binary G4 occupancy and quantitative G4 CUT&Tag signal directly from DNA sequence (Fig. 1). Each input comprised a fixed 1 kb genomic window encoded as a 1,000 × 4 one-hot matrix representing A, C, G and T. The model combined a shared convolutional backbone for local sequence feature extraction with a transformer encoder to capture longer-range dependencies, followed by separate task-specific heads for classification and signal prediction.

Input sequences were processed through two 1D convolutional layers with 64 and 128 filters, respectively, each followed by batch normalization and ReLU activation, to yield base-resolution features, which were average-pooled into 20 sequence tokens (50 bp each), augmented with positional embeddings, and passed through transformer encoder blocks (4-head self-attention, key dimension 32). Each block comprised layer normalization, residual self-attention, and a position-wise feedforward network with GELU activation, with dropout (0.2) applied to the attention and feedforward sublayers.

For binary G4 prediction, transformer tokens were aggregated using an attention-based readout to generate a global sequence representation, which was passed through layer normalization, dropout (0.5), and dense layers to produce a sigmoid probability. For signal prediction, CNN features were mean-pooled to 5 bp resolution (200 bins per 1 kb window), and transformer token embeddings were upsampled to the same resolution. Binned CNN features and upsampled transformer embeddings were then combined with a broadcast global context vector derived from the classification representation and passed through dense layers with dropout (0.3) to predict a 200-bin CUT&Tag signal profile via a linear output layer.

Models were trained separately for each genotype using its corresponding CUT&Tag data. For each 1 kb window, target signal values were extracted from the corresponding bigWig track, mean-pooled into non-overlapping 5 bp bins, transformed using log(1 + x), and globally standardized by z-scoring with parameters estimated on the training split and applied unchanged to validation and test data. To prevent chromosomal leakage, data were partitioned by chromosome, with chr17– 18 used for validation, chr19–22 and chrX for testing, and all remaining chromosomes for training. As a robustness analysis, classification was also repeated across three additional predefined whole-chromosome train, validation and test splits while keeping the pipeline unchanged; split definitions and per-partition results are provided in Supplementary Tables 2 and 3.

Training jointly optimized classification and signal prediction using a weighted multi-task objective,

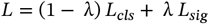

where *L*_*cls*_ was binary cross-entropy, *L*_*sig*_ was Huber loss (δ=1.0), and λ = 0.6 by default. Huber loss was applied to both positive and background windows, enabling the model to learn enriched signal profiles at positive loci and low or flat profiles at negative loci. Models were optimized with Adam (initial learning rate 1 × 10^!”^, gradient clipping (maximum norm 1.0), and batch size 32. Early stopping and model selection were based on validation AUPRC, and the learning rate was reduced on plateau by a factor of 0.5 to a minimum of 1 × 10^!#^ .

For classification, the decision threshold was selected on the validation set by maximizing F1 over thresholds from 0 to 1 in increments of 0.001 and then applied to the held-out test set. Performance was assessed using accuracy, F1, AUROC, and AUPRC for classification, and MAE, MSE, and Pearson correlation on the log-transformed scale for signal prediction. Models were implemented in TensorFlow/Keras and trained on a system equipped with four 18-core Intel Xeon E7-8867 v4 processors (144 logical CPUs) and 1.5 TiB of RAM, using a random seed of 1337. A single model trained in approximately 30–40 min; the genotype–background sweep, chromosome-partition analysis and architectural ablations (50 runs total) required tens of CPU-hours without GPU acceleration.

### Ablation studies

To assess the contribution of individual architectural components, we trained six model variants: (1) the full model described above; (2) without transformer (CNN-only); (3) without signal task (classification only); (4) 1 CNN layer; (5) 1 attention layer; and (6) 50 nodes. Each configuration was trained using the same optimization scheme across multiple random seeds and background definitions.

### Benchmarking of baseline methods

We benchmarked our model against published G4 prediction approaches using the same genotype-specific datasets and chromosome-holdout evaluation framework, restricting comparisons to the most stringent background set, in vitro only GC-matched. Benchmarked methods included three rule-based predictors, Quadparser, G4Hunter, and pqsfinder, four learnable classifiers, G4Beacon, G4detector, G4-Attention, and DeepG4, and one transfer baseline, G4mismatch. Rule-based methods were applied using their published definitions or available software packages. For trainable baselines, public implementations were used or adapted where available; otherwise, models were implemented according to the published architecture. All trainable baselines were trained from scratch on the same genotype-specific CUT&Tag-defined datasets. G4mismatch was instead trained on G4-seq-derived labels to assess transfer from in vitro G4-seq supervision to cell-based CUT&Tag supervision. Additional implementation details and model complexity summaries are provided in Supplementary Tables 9 and 14. All learnable baselines except G4mismatch were trained on chr1-16, validated on chr17-18, and evaluated on held-out test chromosomes chr19-22 and chrX. For all methods except G4mismatch, thresholds were selected using the same validation-set F1-maximization procedure described above. AUROC and AUPRC were computed from continuous test-set scores, and F1 was computed at the validation-selected threshold. G4mismatch was evaluated only as a zero-shot transfer baseline, so only AUROC and AUPRC are reported. Signal regression was not benchmarked because, unlike our model, the comparison methods were not trained to predict CUT&Tag signal intensity.

### Independent application and evaluation in K562 cells

Independent application of the G4SCOPE framework used published K562 BG4 CUT&Tag data (GEO accession GSE181373). BG4 peaks were centered and expanded to 1 kb windows (n = 18,357) after excluding windows extending beyond chromosome boundaries or containing ambiguous bases. The same negative-set definitions as the primary analysis were applied (random, GC-matched, in vitro only, and in vitro only GC-matched), and the same chromosome split was used (train chr1–16; validation chr17–18; test chr19–22 plus chrX). The same architecture described above was

### G4 subtyping via learned sequence embeddings and genomic context

To identify subclasses of G-quadruplex (G4) loci, we clustered model-derived sequence embeddings from G4-positive loci across genotypes. Attention-weighted transformer embeddings from the 1 kb input sequences were standardized, reduced by principal component analysis (PCA) to 30 components accounting for >99.9% of the variance, and clustered using Gaussian mixture models (GMMs) with full covariance matrices. Models with 2–10 components were compared using the Bayesian Information Criterion (BIC), and loci were assigned to clusters by maximum posterior probability. UMAP was used only for visualization and did not influence cluster assignment. To biologically interpret the resulting clusters, we examined directly interpretable sequence and genomic features, including GC content, CpG density, promoters, TSSs, 1–5 kb TSS-flanking regions, CpG islands, shores and shelves, exons, introns, intron–exon boundaries, intergenic regions, enhancers, 5′ and 3′ UTRs, and distance to CTCF peaks. GC content and CpG density were calculated from the full 1 kb input sequence, with ambiguous bases excluded. CTCF proximity was categorized as ≤1 kb, 1–5 kb, 5–10 kb or >10 kb. Categorical annotations were tested using Fisher’s exact test and summarized as log2 odds ratios, whereas continuous features were compared using the Mann–Whitney test and summarized as log2 median ratios. P values were adjusted using the Benjamini–Hochberg procedure, and associations with FDR < 0.05 were retained. Significant effects were standardized within each feature across clusters for visualization; undefined median-ratio effects were treated as missing. WT clusters with similar enrichment profiles were further grouped into five meta-clusters: Regulatory/CpG-rich, CTCF-proximal, CpG-poor promoter-depleted/CTCF-neutral, CpG-poor/CTCF-distal and Mixed/weak. Meta-cluster effects were calculated as the median effect across constituent clusters and standardized across the five meta-clusters.

