## Supplementary material for "G4SCOPE identifies sequence features and epigenetic contexts of chromatin G-quadruplexes"

---

### Contents

#### Tables

**Supplementary Table 1. Classification performance across genotypes and negative sets**
**Supplementary Table 2. Chromosome split definitions used for chromosome-partition robustness analysis**
**Supplementary Table 3. Classification performance across chromosome partitions on the in vitro only GC-matched benchmark**
**Supplementary Table 4. Regression performance across genotypes and negative sets**
**Supplementary Table 5. Ablation study for the WT genotype**
**Supplementary Table 6. Ablation study for the TP53−/− genotype**
**Supplementary Table 7. Ablation study for the TP53−/−;BRCA1−/− genotype**
**Supplementary Table 8. Ablation study for the TP53−/−;BRCA2−/− genotype**
**Supplementary Table 9. Implementation source for each benchmarked method**
**Supplementary Table 10. Overall benchmark classification performance of G4 prediction methods**
**Supplementary Table 11. Genotype-level benchmark AUROC on the in vitro only GC-matched benchmark**
**Supplementary Table 12. Genotype-level benchmark AUPRC on the in vitro only GC-matched benchmark**
**Supplementary Table 13. Genotype-level benchmark F1 on the in vitro only GC-matched benchmark**
**Supplementary Table 14. Model complexity and inference time**
**Supplementary Table 15. K562 external validation**
**Supplementary Table 16. Benchmark comparison of G4 prediction methods in K562 cells**

#### Figures

**Supplementary Figure 1. Confusion matrices for binary G4 classification across all 16 experimental conditions**
**Supplementary Figure 2. Calibration curves for the G4 classification model across all 16 experimental conditions**
**Supplementary Figure 3. Robustness of learned G4 sequence motifs across negative training-set definitions**
**Supplementary Figure 4. Cross-genotype transfer of multi-task G4 models**
**Supplementary Figure 5. Ablation analysis of the multi-task convolutional–transformer model across genotypes**
**Supplementary Figure 6. Gaussian mixture model selection for G4 embedding clusters**
**Supplementary Figure 7. Localisation of transcription-factor family motif enrichment across predicted clusters**
**Supplementary Figure 8. Positional distributions of transcription-factor family motifs across predicted WT clusters.**

**Supplementary Table 1. Classification performance across genotypes and negative sets.** Mean ± SD across 3 seeds for each genotype × negative-set combination.

| Genotype | Neg. | AUROC | AUPRC | F1 | Acc. |
|---|---|---|---|---|---|
| WT | random | 0.955 ± 0.016 | 0.952 ± 0.015 | 0.883 ± 0.021 | 0.877 ± 0.022 |
| | In vitro only | 0.946 ± 0.018 | 0.947 ± 0.017 | 0.871 ± 0.023 | 0.866 ± 0.024 |
| | In vitro only, GC-matched | 0.906 ± 0.022 | 0.908 ± 0.021 | 0.833 ± 0.026 | 0.831 ± 0.027 |
| | GC-matched | 0.907 ± 0.023 | 0.902 ± 0.022 | 0.833 ± 0.027 | 0.830 ± 0.028 |
| TP53−/− | random | 0.965 ± 0.014 | 0.964 ± 0.014 | 0.899 ± 0.018 | 0.897 ± 0.019 |
| | In vitro only | 0.948 ± 0.017 | 0.948 ± 0.017 | 0.864 ± 0.021 | 0.853 ± 0.022 |
| | In vitro only, GC-matched | 0.914 ± 0.021 | 0.914 ± 0.020 | 0.843 ± 0.024 | 0.841 ± 0.025 |
| | GC-matched | 0.910 ± 0.022 | 0.907 ± 0.021 | 0.832 ± 0.026 | 0.831 ± 0.027 |
| TP53−/−;BRCA1−/− | random | 0.953 ± 0.017 | 0.951 ± 0.016 | 0.880 ± 0.020 | 0.874 ± 0.021 |
| | In vitro only | 0.938 ± 0.019 | 0.940 ± 0.018 | 0.855 ± 0.022 | 0.848 ± 0.023 |
| | In vitro only, GC-matched | 0.900 ± 0.023 | 0.903 ± 0.022 | 0.822 ± 0.025 | 0.819 ± 0.026 |
| | GC-matched | 0.889 ± 0.024 | 0.888 ± 0.023 | 0.807 ± 0.027 | 0.801 ± 0.028 |
| TP53−/−;BRCA2−/− | random | 0.963 ± 0.015 | 0.962 ± 0.015 | 0.892 ± 0.019 | 0.887 ± 0.020 |
| | In vitro only | 0.951 ± 0.018 | 0.945 ± 0.018 | 0.879 ± 0.022 | 0.877 ± 0.023 |
| | In vitro only, GC-matched | 0.927 ± 0.020 | 0.926 ± 0.020 | 0.849 ± 0.024 | 0.846 ± 0.025 |
| | GC-matched | 0.908 ± 0.022 | 0.909 ± 0.021 | 0.836 ± 0.026 | 0.827 ± 0.027 |

**Supplementary Table 2. Chromosome split definitions used for chromosome-partition robustness analysis.** Each split used two validation chromosomes and five test chromosomes.

| Split ID | Validation chromosomes | Test chromosomes |
|---|---|---|
| primary | chr17, chr18 | chr19, chr20, chr21, chr22, chrX |
| alt1_mixed | chr3, chr15 | chr4, chr8, chr11, chr16, chr20 |
| alt2_mixed | chr6, chr13 | chr1, chr9, chr14, chr21, chrX |
| alt3_mixed | chr5, chr19 | chr2, chr10, chr12, chr18, chr22 |

**Supplementary Table 3. Classification performance across chromosome partitions on the In vitro only GC-matched benchmark.**

| Chromosome split | Genotype | AUROC | AUPRC | F1 | Accuracy | Epochs |
|---|---|---|---|---|---|---|
| alt1_mixed | WT | 0.8978 | 0.8929 | 0.8229 | 0.8154 | 10 |
| alt1_mixed | TP53−/− | 0.9084 | 0.9135 | 0.8358 | 0.8337 | 14 |
| alt1_mixed | TP53−/−;BRCA1−/− | 0.8909 | 0.9008 | 0.8189 | 0.8192 | 12 |
| alt1_mixed | TP53−/−;BRCA2−/− | 0.9342 | 0.9318 | 0.8617 | 0.8624 | 9 |
| alt2_mixed | WT | 0.9027 | 0.9021 | 0.8318 | 0.8291 | 14 |
| alt2_mixed | TP53−/− | 0.8958 | 0.8839 | 0.8264 | 0.8247 | 9 |
| alt2_mixed | TP53−/−;BRCA1−/− | 0.8886 | 0.8837 | 0.8056 | 0.8124 | 9 |
| alt2_mixed | TP53−/−;BRCA2−/− | 0.9128 | 0.9048 | 0.8401 | 0.8389 | 9 |
| alt3_mixed | WT | 0.8957 | 0.8941 | 0.8162 | 0.8111 | 13 |
| alt3_mixed | TP53−/− | 0.8931 | 0.8894 | 0.8199 | 0.8048 | 12 |
| alt3_mixed | TP53−/−;BRCA1−/− | 0.8846 | 0.8843 | 0.8136 | 0.7988 | 12 |
| alt3_mixed | TP53−/−;BRCA2−/− | 0.9209 | 0.9174 | 0.8487 | 0.8365 | 11 |

**Supplementary Table 4. Regression performance across genotypes and negative sets.** Mean ± SD across 3 seeds for each genotype × negative-set combination.

| Genotype | Neg. | MAE | MSE | r |
|---|---|---|---|---|
| WT | random | 0.140 ± 0.011 | 0.067 ± 0.006 | 0.70 ± 0.025 |
| | In vitro only | 0.146 ± 0.012 | 0.065 ± 0.007 | 0.70 ± 0.026 |
| | In vitro only, GC-matched | 0.163 ± 0.015 | 0.079 ± 0.009 | 0.62 ± 0.028 |
| | GC-matched | 0.172 ± 0.016 | 0.076 ± 0.009 | 0.64 ± 0.029 |
| TP53−/− | random | 0.133 ± 0.011 | 0.054 ± 0.005 | 0.71 ± 0.026 |
| | In vitro only | 0.134 ± 0.011 | 0.055 ± 0.006 | 0.70 ± 0.027 |
| | In vitro only, GC-matched | 0.150 ± 0.014 | 0.060 ± 0.007 | 0.68 ± 0.028 |
| | GC-matched | 0.149 ± 0.014 | 0.064 ± 0.007 | 0.65 ± 0.029 |
| TP53−/−;BRCA1−/− | random | 0.113 ± 0.009 | 0.040 ± 0.004 | 0.70 ± 0.024 |
| | In vitro only | 0.118 ± 0.010 | 0.042 ± 0.005 | 0.67 ± 0.025 |
| | In vitro only, GC-matched | 0.128 ± 0.012 | 0.045 ± 0.005 | 0.64 ± 0.027 |
| | GC-matched | 0.129 ± 0.013 | 0.048 ± 0.006 | 0.62 ± 0.028 |
| TP53−/−;BRCA2−/− | random | 0.085 ± 0.007 | 0.026 ± 0.003 | 0.72 ± 0.030 |
| | In vitro only | 0.099 ± 0.009 | 0.031 ± 0.004 | 0.65 ± 0.028 |
| | In vitro only, GC-matched | 0.092 ± 0.009 | 0.032 ± 0.004 | 0.67 ± 0.027 |
| | GC-matched | 0.093 ± 0.010 | 0.029 ± 0.004 | 0.68 ± 0.028 |

**Supplementary Table 5. Ablation study, WT genotype.** Mean ± SD across 3 seeds × 4 negative sets (n = 12 runs per row).

| Configuration | Params | AUROC | AUPRC | F1 | Accuracy | Signal r |
|---|---|---|---|---|---|---|
| Full model | 178K | 0.929 ± 0.024 | 0.926 ± 0.023 | 0.854 ± 0.027 | 0.851 ± 0.027 | 0.670 ± 0.027 |
| w/o Transformer | 57K | 0.808 ± 0.029 | 0.809 ± 0.030 | 0.728 ± 0.029 | 0.722 ± 0.027 | 0.485 ± 0.028 |
| w/o Signal task | 149K | 0.861 ± 0.028 | 0.859 ± 0.028 | 0.785 ± 0.031 | 0.780 ± 0.032 | — |
| 1 CNN layer | 149K | 0.840 ± 0.027 | 0.849 ± 0.027 | 0.772 ± 0.026 | 0.765 ± 0.028 | 0.574 ± 0.030 |
| 1 Attn layer | 128K | 0.868 ± 0.024 | 0.864 ± 0.023 | 0.794 ± 0.025 | 0.788 ± 0.026 | 0.597 ± 0.033 |
| 50 nodes | 178K | 0.859 ± 0.025 | 0.855 ± 0.026 | 0.783 ± 0.026 | 0.778 ± 0.026 | 0.572 ± 0.036 |

**Supplementary Table 6. Ablation study, TP53−/− genotype.** Mean ± SD across 3 seeds × 4 negative sets (n = 12 runs per row).

| Configuration | Params | AUROC | AUPRC | F1 | Accuracy | Signal r |
|---|---|---|---|---|---|---|
| Full model | 178K | 0.932 ± 0.015 | 0.932 ± 0.015 | 0.854 ± 0.015 | 0.851 ± 0.014 | 0.682 ± 0.019 |
| w/o Transformer | 57K | 0.819 ± 0.018 | 0.821 ± 0.021 | 0.737 ± 0.013 | 0.729 ± 0.013 | 0.499 ± 0.022 |
| w/o Signal task | 149K | 0.864 ± 0.021 | 0.860 ± 0.021 | 0.788 ± 0.019 | 0.781 ± 0.020 | — |
| 1 CNN layer | 149K | 0.843 ± 0.027 | 0.853 ± 0.028 | 0.774 ± 0.022 | 0.766 ± 0.019 | 0.594 ± 0.028 |
| 1 Attn layer | 128K | 0.870 ± 0.016 | 0.868 ± 0.016 | 0.795 ± 0.019 | 0.792 ± 0.017 | 0.627 ± 0.027 |
| 50 nodes | 178K | 0.864 ± 0.016 | 0.866 ± 0.017 | 0.784 ± 0.013 | 0.779 ± 0.012 | 0.587 ± 0.017 |

**Supplementary Table 7. Ablation study, TP53−/−;BRCA1−/− genotype.** Mean ± SD across 3 seeds × 4 negative sets (n = 12 runs per row)

| Configuration | Params | AUROC | AUPRC | F1 | Accuracy | Signal r |
|---|---|---|---|---|---|---|
| Full model | 178K | 0.918 ± 0.026 | 0.918 ± 0.024 | 0.843 ± 0.020 | 0.838 ± 0.018 | 0.659 ± 0.014 |
| w/o Transformer | 57K | 0.801 ± 0.019 | 0.806 ± 0.018 | 0.717 ± 0.014 | 0.708 ± 0.012 | 0.486 ± 0.020 |
| w/o Signal task | 149K | 0.849 ± 0.023 | 0.850 ± 0.019 | 0.768 ± 0.025 | 0.762 ± 0.023 | — |
| 1 CNN layer | 149K | 0.830 ± 0.023 | 0.842 ± 0.023 | 0.755 ± 0.024 | 0.751 ± 0.016 | 0.571 ± 0.033 |
| 1 Attn layer | 128K | 0.857 ± 0.014 | 0.857 ± 0.013 | 0.776 ± 0.011 | 0.770 ± 0.007 | 0.598 ± 0.016 |
| 50 nodes | 178K | 0.849 ± 0.023 | 0.849 ± 0.024 | 0.772 ± 0.021 | 0.766 ± 0.016 | 0.567 ± 0.035 |

**Supplementary Table 8. Ablation study, TP53−/−;BRCA2−/− genotype.** Mean ± SD across 3 seeds × 4 negative sets (n = 12 runs per row)

| Configuration | Params | AUROC | AUPRC | F1 | Accuracy | Signal r |
|---|---|---|---|---|---|---|
| Full model | 178K | 0.937 ± 0.024 | 0.935 ± 0.026 | 0.862 ± 0.026 | 0.858 ± 0.027 | 0.688 ± 0.039 |
| w/o Transformer | 57K | 0.813 ± 0.028 | 0.817 ± 0.029 | 0.731 ± 0.026 | 0.730 ± 0.028 | 0.483 ± 0.026 |
| w/o Signal task | 149K | 0.870 ± 0.029 | 0.868 ± 0.030 | 0.796 ± 0.031 | 0.792 ± 0.035 | — |
| 1 CNN layer | 149K | 0.847 ± 0.030 | 0.854 ± 0.031 | 0.781 ± 0.034 | 0.778 ± 0.034 | 0.590 ± 0.029 |
| 1 Attn layer | 128K | 0.877 ± 0.022 | 0.872 ± 0.024 | 0.804 ± 0.028 | 0.802 ± 0.029 | 0.629 ± 0.035 |
| 50 nodes | 178K | 0.868 ± 0.028 | 0.865 ± 0.030 | 0.794 ± 0.031 | 0.789 ± 0.032 | 0.569 ± 0.039 |

**Supplementary Table 9. Implementation source for each benchmarked method.** Source and implementation mode for each benchmarked method. Rule-based methods were coded from published definitions or run with the cited package. Learnable baselines were trained on the same genotype-specific CUT&Tag datasets and chromosome partitions as the main model. G4mismatch was trained on In vitro only labels and evaluated on CUT&Tag without CUT&Tag-specific fine-tuning.

| Method | Implementation source | Notes |
|---|---|---|
| Quadparser | Implemented from Huppert & Balasubramanian (2005) | No official software available; canonical PQS regular expression used on both strands; per-window motif density (quadparser_density) used as score |
| G4Hunter | Implemented from Bedrat et al. (2016) | Published G-richness / G-skewness algorithm with a 25 bp sliding window; maximum absolute window score (g4hunter_max) used as region score |
| pqsfinder | Used Bioconductor R package pqsfinder (Hon et al., 2017) | Installed from Bioconductor; maximum pqsfinder score per window (pqsfinder_max) used as region score |
| G4Beacon | Re-implemented from Zhang et al. (2023) | Gradient-boosting (LightGBM) re-implemented for this benchmark using sequence-only input (one-hot encoded 1 kb windows, no ATAC-seq), for fair comparison with other sequence-based baselines |
| G4detector | Implemented from Barshai et al. (2022) | CNN architecture reproduced from the paper; trained on our data with the same chromosome holdout; no official code or pre-trained weights used |
| G4-Attention | Implemented from Mukherjee et al. (2024) | CNN + BiLSTM + attention classification architecture reproduced; the original study's regression head for In vitro only mismatch scores was not used, as it is not comparable to CUT&Tag signal regression |
| DeepG4 | Re-implemented from Rocher et al. (2021) | Multi-scale CNN re-implemented and trained in our pipeline with identical inputs and chromosome splits |
| G4mismatch | Implemented from Barshai et al. (2023) | CNN implemented as a zero-shot transfer baseline; trained on In vitro only-derived labels and evaluated on CUT&Tag without CUT&Tag-specific fine-tuning or threshold selection |

**Supplementary Table 10. Overall benchmark classification performance of G4 prediction methods.** Values are mean ± SD across four genotypes (WT, TP53−/−, TP53−/−;BRCA1−/−, and TP53−/−;BRCA2−/−). AUROC and AUPRC were computed from continuous test-set scores. F1 was computed using validation-selected thresholds for methods evaluated with CUT&Tag threshold calibration. G4mismatch was evaluated as a zero-shot transfer baseline trained on In vitro only labels, so F1 is not reported.

| **Method** | **AUROC** | **AUPRC** | **F1** |
|---|---|---|---|
| Our model | 0.912 ± 0.012 | 0.913 ± 0.010 | 0.837 ± 0.012 |
| DeepG4 | 0.881 ± 0.024 | 0.862 ± 0.024 | 0.801 ± 0.016 |
| G4-Attention | 0.877 ± 0.029 | 0.856 ± 0.048 | 0.827 ± 0.013 |
| G4detector | 0.771 ± 0.050 | 0.793 ± 0.031 | 0.748 ± 0.016 |
| G4Beacon | 0.654 ± 0.005 | 0.657 ± 0.012 | 0.666 ± 0.001 |
| pqsfinder | 0.509 ± 0.003 | 0.517 ± 0.003 | 0.500 ± 0.010 |
| Quadparser | 0.505 ± 0.003 | 0.500 ± 0.002 | 0.500 ± 0.000 |
| G4Hunter | 0.488 ± 0.004 | 0.493 ± 0.002 | 0.490 ± 0.000 |
| G4mismatch | 0.202 ± 0.052 | 0.353 ± 0.030 | — |

**Supplementary Table 11. Genotype-level benchmark AUROC on the In vitro only GC-matched benchmark**. AUROC values for each benchmarked method across the four genotypes.

| Method | WT | TP53−/− | TP53−/−;BRCA1−/− | TP53−/−;BRCA2−/− |
|---|---|---|---|---|
| Our model | 0.906 ± 0.022 | 0.914 ± 0.021 | 0.900 ± 0.023 | 0.927 ± 0.020 |
| DeepG4 | 0.903 | 0.880 | 0.847 | 0.892 |
| G4-Attention | 0.869 | 0.871 | 0.850 | 0.916 |
| G4detector | 0.727 | 0.729 | 0.789 | 0.838 |
| G4Beacon | 0.649 | 0.656 | 0.652 | 0.659 |
| pqsfinder | 0.506 | 0.507 | 0.512 | 0.511 |
| Quadparser | 0.503 | 0.503 | 0.509 | 0.505 |
| G4Hunter | 0.488 | 0.483 | 0.492 | 0.487 |
| G4mismatch | 0.276 | 0.179 | 0.182 | 0.170 |

**Supplementary Table 12. Genotype-level benchmark AUPRC on the In vitro only GC-matched benchmark.** AUPRC values for each benchmarked method across the four genotypes.

| Method | WT | TP53−/− | TP53−/−;BRCA1−/− | TP53−/−;BRCA2−/− |
|---|---|---|---|---|
| Our model | 0.908 ± 0.021 | 0.914 ± 0.020 | 0.903 ± 0.022 | 0.926 ± 0.020 |
| DeepG4 | 0.874 | 0.850 | 0.835 | 0.890 |
| G4-Attention | 0.853 | 0.798 | 0.862 | 0.912 |
| G4detector | 0.778 | 0.792 | 0.765 | 0.838 |
| G4Beacon | 0.644 | 0.663 | 0.646 | 0.671 |
| pqsfinder | 0.512 | 0.516 | 0.519 | 0.518 |
| Quadparser | 0.499 | 0.499 | 0.504 | 0.499 |
| G4Hunter | 0.492 | 0.494 | 0.494 | 0.491 |
| G4mismatch | 0.399 | 0.338 | 0.338 | 0.335 |

**Supplementary Table 13. Genotype-level benchmark F1 on the In vitro only GC-matched benchmark.** F1 values for each benchmarked method across the four genotypes. F1 was computed on the held-out test set using the threshold that maximized validation-set F1 for methods evaluated with CUT&Tag threshold calibration. G4mismatch was evaluated as a zero-shot transfer baseline trained on In vitro only labels, without CUT&Tag threshold selection, so F1 is not reported.

| Method | WT | TP53−/− | TP53−/−;BRCA1−/− | TP53−/−;BRCA2−/− |
|---|---|---|---|---|
| Our model | 0.833 ± 0.026 | 0.843 ± 0.024 | 0.822 ± 0.025 | 0.849 ± 0.024 |
| DeepG4 | 0.800 | 0.806 | 0.780 | 0.818 |
| G4-Attention | 0.828 | 0.831 | 0.809 | 0.839 |
| G4detector | 0.763 | 0.733 | 0.733 | 0.762 |
| G4Beacon | 0.666 | 0.667 | 0.666 | 0.664 |
| pqsfinder | 0.500 | 0.500 | 0.510 | 0.510 |
| Quadparser | 0.500 | 0.500 | 0.500 | 0.500 |
| G4Hunter | 0.490 | 0.490 | 0.490 | 0.490 |
| G4mismatch | — | — | — | — |

**Supplementary Table 14. Model complexity and inference time.** Parameter counts are exact values logged during training. Inference time is reported per sequence for 1 kb one-hot encoded input. Complexity class refers to the dominant per-sequence computational complexity as a function of sequence length L. For G4-Attention, only the classification output was evaluated, as its auxiliary regression target, In vitro only mismatch score, is not comparable to CUT&Tag signal intensity. Dashes indicate non-neural methods for which parameter count is not applicable.

| Method | Architecture | Parameters | Tasks | Inference complexity | Inference time (ms/seq) | Training required |
|---|---|---|---|---|---|---|
| Our model | Multi-scale CNN + Transformer (self-attention) | 178K | Classification + CUT&Tag signal regression (in vivo) | $O(L^2)$ | 3–8 | Yes |
| G4-Attention | CNN + BiLSTM + attention | 118K | Classification + In vitro only mismatch regression (in vitro) | $O(L^2)$ | 3–8 | Yes |
| G4detector | CNN | 107K | Classification only | $O(L)$ | 0.5–2 | Yes |
| G4mismatch | CNN | 107K | Classification only (transfer) | $O(L)$ | 0.5–2 | Yes |
| DeepG4 | Multi-scale CNN | 60K | Classification only | $O(L)$ | 0.5–2 | Yes |
| G4Beacon | LightGBM | — | Classification only | $O(L)$ | 1–3 | Yes |
| pqsfinder | PQS search | — | Classification only | $O(L^2)$ | 10–30 | No |
| G4Hunter | Sliding-window score | — | Classification only | $O(L)$ | 0.1–0.5 | No |
| Quadparser | Regex scan | — | Classification only | $O(L)$ | 0.1–0.5 | No |

**Supplementary Table 15: K562 external validation.** Performance of the multi-task model trained from scratch on K562 BG4 CUT&Tag peaks (GSE181373) using the primary chromosome-holdout split (train chr1–16; validation chr17–18; test chr19–22 plus chrX). Metrics are reported on held-out test chromosomes; signal *r* is Pearson correlation between predicted and observed 5 bp-binned BG4 CUT&Tag signal.

| Negative set | AUROC | AUPRC | F1 | Accuracy | Signal *r* |
|---|---|---|---|---|---|
| GC-matched | 0.918 | 0.907 | 0.841 | 0.836 | 0.618 |
| In vitro only, GC-matched | 0.924 | 0.916 | 0.849 | 0.844 | 0.623 |
| In vitro only | 0.941 | 0.934 | 0.869 | 0.862 | 0.641 |
| Random | 0.953 | 0.947 | 0.882 | 0.876 | 0.658 |

**Supplementary Table 16. Benchmark comparison of G4 prediction methods in K562 cells.** Benchmark performance on the K562 GC-matched BG4 CUT&Tag evaluation (GSE181373). AUROC and AUPRC were computed from continuous test-set scores. F1 was computed using validation-selected thresholds for methods trained on CUT&Tag labels; G4mismatch was evaluated as a In vitro only transfer baseline, so F1 is not reported.

| Method | AUROC | AUPRC | F1 |
|---|---|---|---|
| Our model | 0.918 | 0.907 | 0.841 |
| DeepG4 | 0.874 | 0.869 | 0.793 |
| G4-Attention | 0.883 | 0.849 | 0.831 |
| G4detector | 0.764 | 0.798 | 0.752 |
| G4Beacon | 0.661 | 0.651 | 0.659 |
| pqsfinder | 0.510 | 0.520 | 0.500 |
| Quadparser | 0.500 | 0.500 | 0.500 |
| G4Hunter | 0.500 | 0.500 | 0.490 |
| G4mismatch | 0.470 | 0.470 | — |

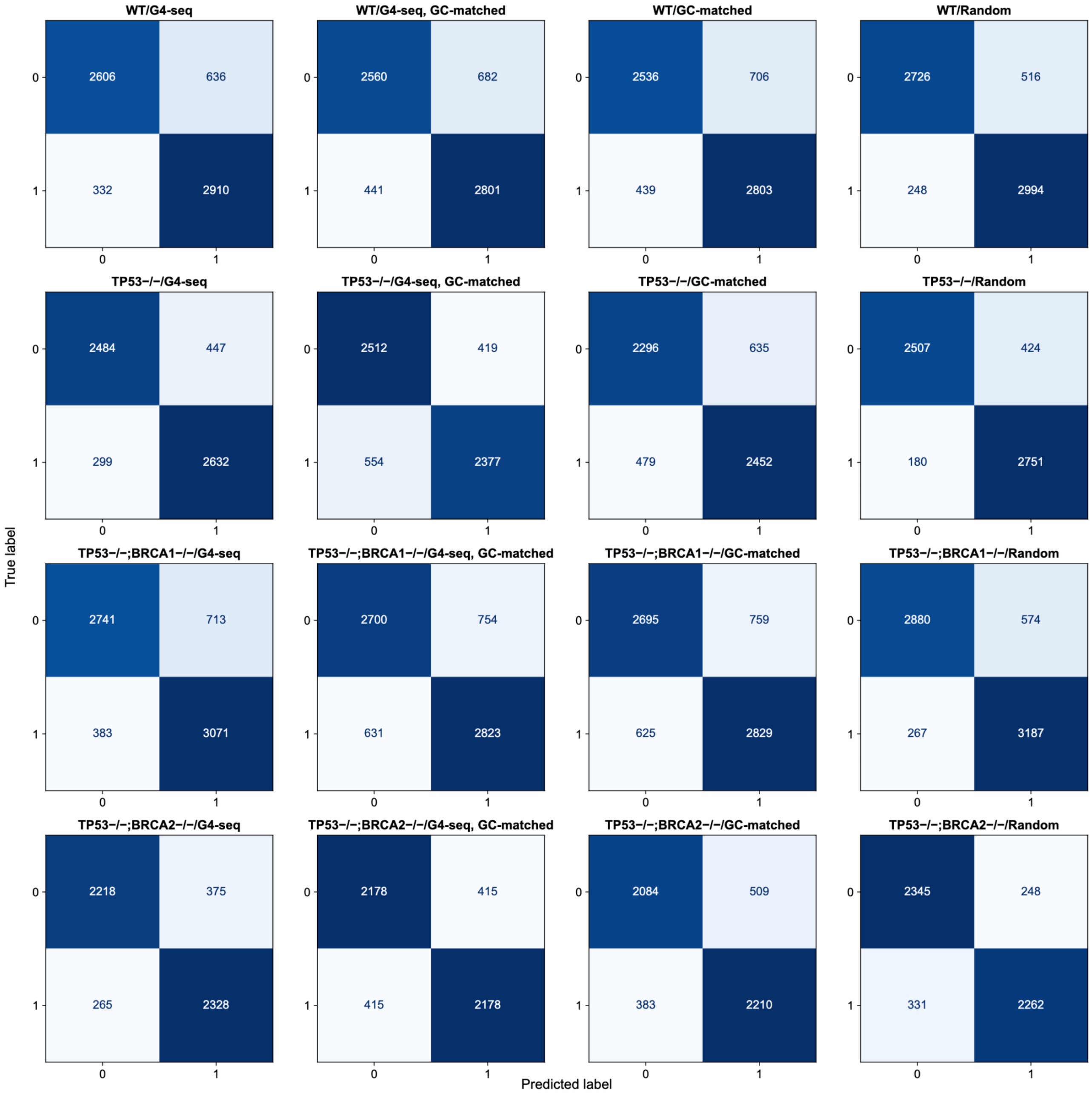


**Supplementary Figure 1. Confusion matrices for binary G4 classification across all 16 experimental conditions.** Confusion matrices showing classification performance of the multi-task convolutional-transformer model for each combination of cell line (rows) and background strategy (columns). Rows indicate the true label and columns the predicted label, where 0 = non-G4 and 1 = G4. Cell counts reflect the number of test-set windows assigned to each category; color intensity is proportional to count magnitude. High true-positive and true-negative counts relative to misclassifications are consistent across all conditions.

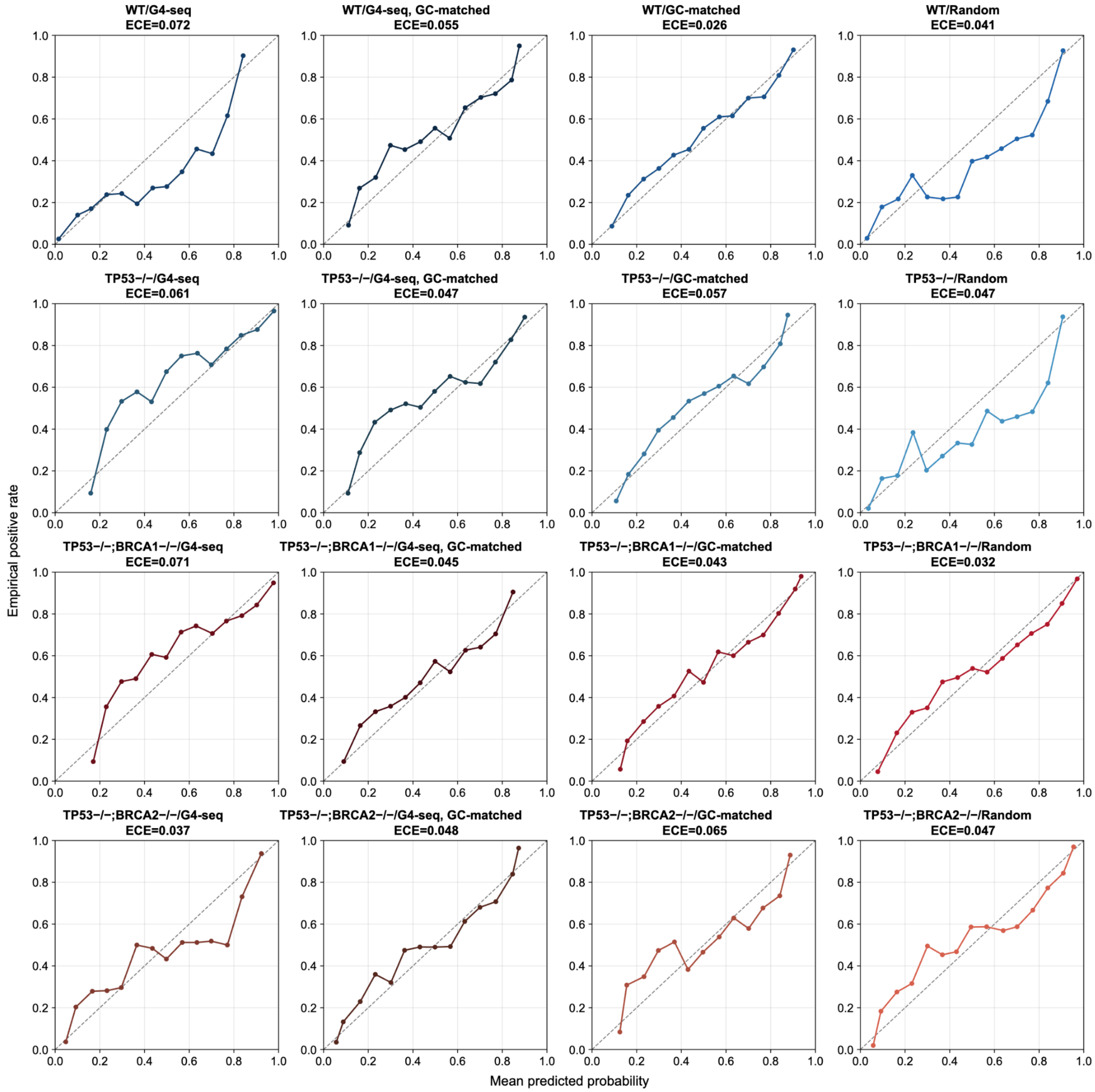


**Supplementary Figure 2. Calibration curves for the G4 classification model across all 16 experimental conditions.** Reliability diagrams assessing the calibration of predicted probabilities for each combination of cell line (rows) and negative-set strategy (columns). Each panel plots the empirical positive rate against the mean predicted probability across equal-frequency bins; the dashed diagonal represents perfect calibration. Expected Calibration Error (ECE) is shown for each condition (range: 0.026–0.072). Curves closely follow the diagonal across all conditions, indicating that the model's output probabilities are well-calibrated and can be interpreted as reliable estimates of G4-forming likelihood. Panel color reflects cell line identity, consistent with Fig. 3.

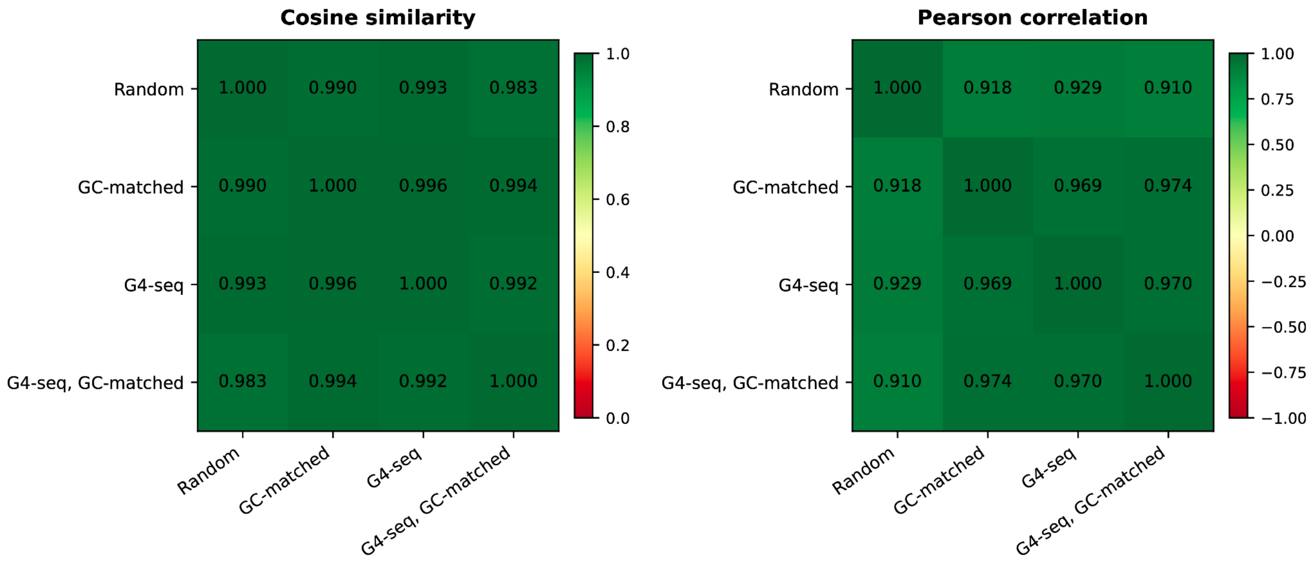


**Supplementary Figure 3. Robustness of learned G4 sequence motifs (Fig. 3E) across negative training set definitions.** Pairwise similarity between saliency-weighted sequence profiles from the WT genotype model trained against each of four negative-set conditions (Random, GC-matched, In vitro only, In vitro only GC-matched), derived from saliency-weighted one-hot sequences of up to 500 positive test examples aggregated into 20 positional bins. (Left) Cosine similarity and (Right) Pearson correlation between flattened PWM vectors. Both metrics exceed 0.98 and 0.91, respectively, across all comparisons, indicating a stable G4-associated motif regardless of negative-set definition. The modest drop in Pearson correlation for the Random set (0.91–0.93 vs. 0.97 for composition-controlled sets) suggests residual GC-content sensitivity in that model, while harder negatives converge on a purer G4 sequence grammar.

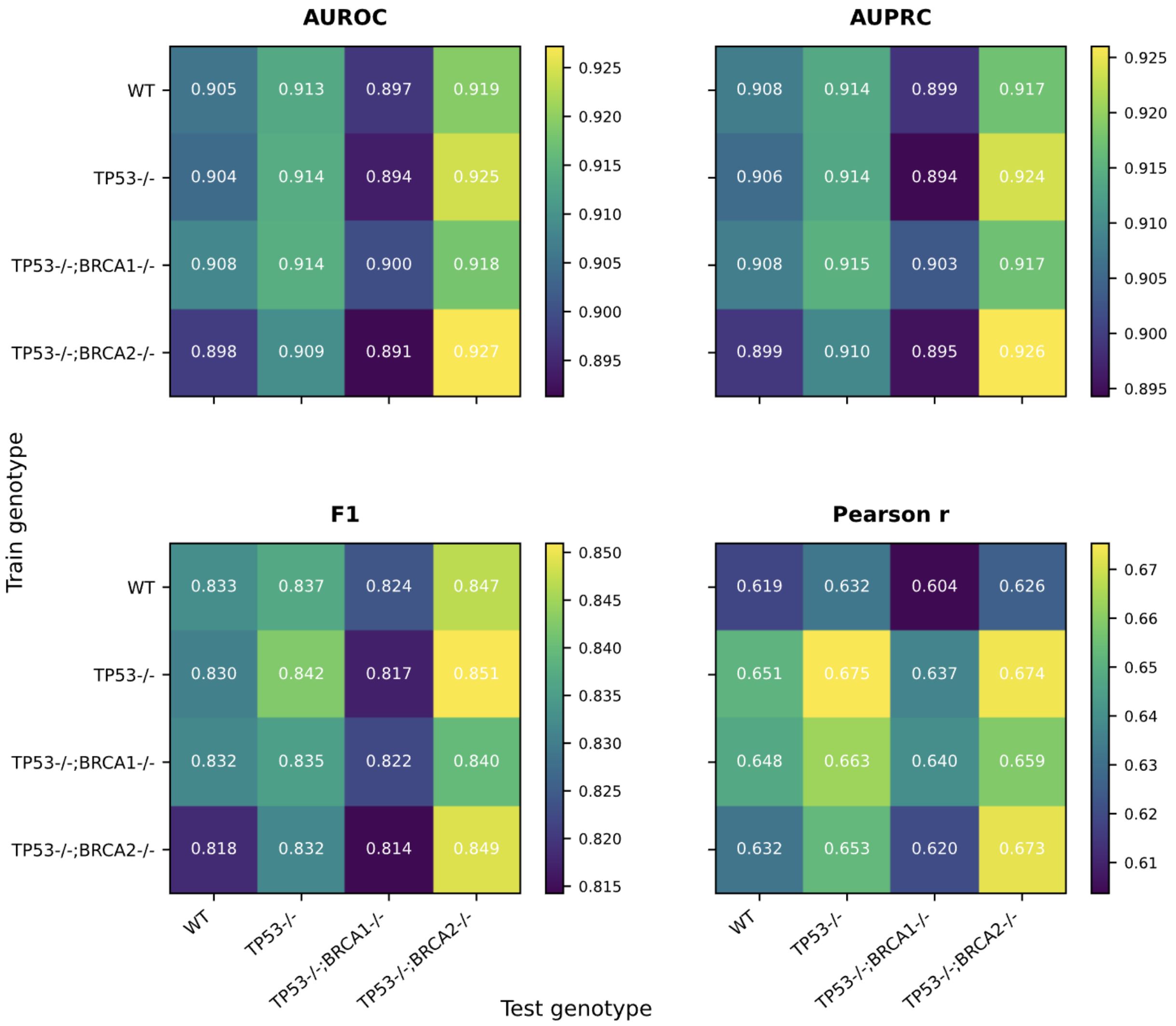


**Supplementary Figure 4. Cross-genotype transfer of multi-task G4 models.** Heatmaps summarize performance when genotype-specific multi-task models trained on one genotype (rows) were evaluated on the held-out test chromosomes of each genotype (columns): WT, TP53−/−, TP53−/−;BRCA1−/−, and TP53−/−;BRCA2−/−, using the In vitro only GC-matched negative set. AUROC and AUPRC are threshold-independent classification metrics. F1 was calculated using the threshold that maximized validation-set F1 in the training genotype and was then applied unchanged to each test genotype. Pearson *r* denotes the correlation between predicted and observed CUT&Tag signal profiles in the test set. Diagonal cells correspond to matched training and test genotypes, whereas off-diagonal cells represent cross-genotype transfer. Performance remained strong across both matched and cross-genotype evaluations, indicating substantial transferability of learned sequence features across genotypes.

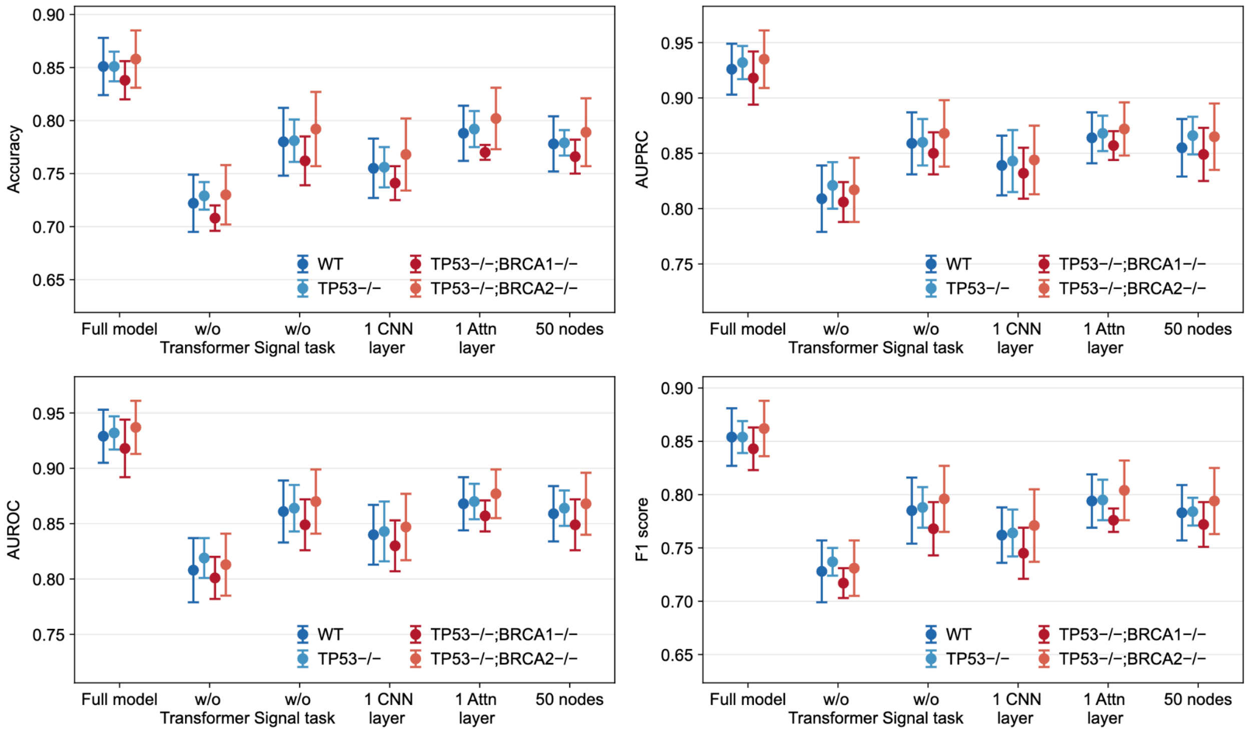


**Supplementary Figure 5. Ablation analysis of the multitask GCNN across genotype classes.** Performance of the multitask GCNN under architectural/task ablations, shown for Accuracy (top-left), AUPRC (top-right), AUROC (bottom-left), and F1 score (bottom-right). We compare the full model against variants with no transformer, no signal task, one CNN layer, one attention layer, and a 50-node sequence representation. Results are reported for four genotype groups: WT, TP53−/−, TP53−/−;BRCA1−/−, and TP53−/−;BRCA2−/−. Points denote model performance, and error bars indicate variability across evaluation runs.

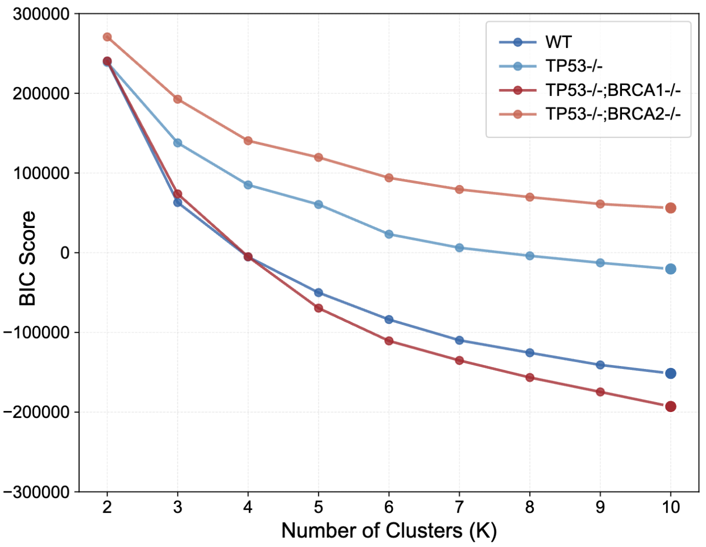


**Supplementary Figure 6. Gaussian mixture model selection for G4 embedding clusters.** Bayesian information criterion (BIC) across Gaussian mixture models with 2–10 components used to select the number of clusters for the G4-trained sequence embeddings.

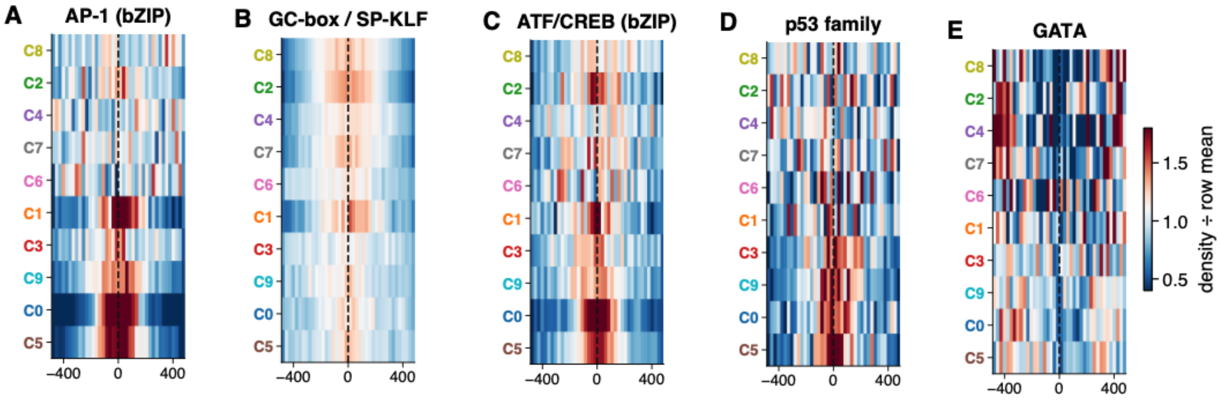


**Supplementary Figure 7.** Examples of localisation of transcription factor motif enrichment across predicted clusters. Heatmaps show positional enrichment patterns for representative TF families, including **(A)** AP-1 (bZIP), **(B)** GC-box/SP-KLF, **(C)** ATF/CREB (bZIP), **(D)** p53 family, and **(E)** GATA motifs.

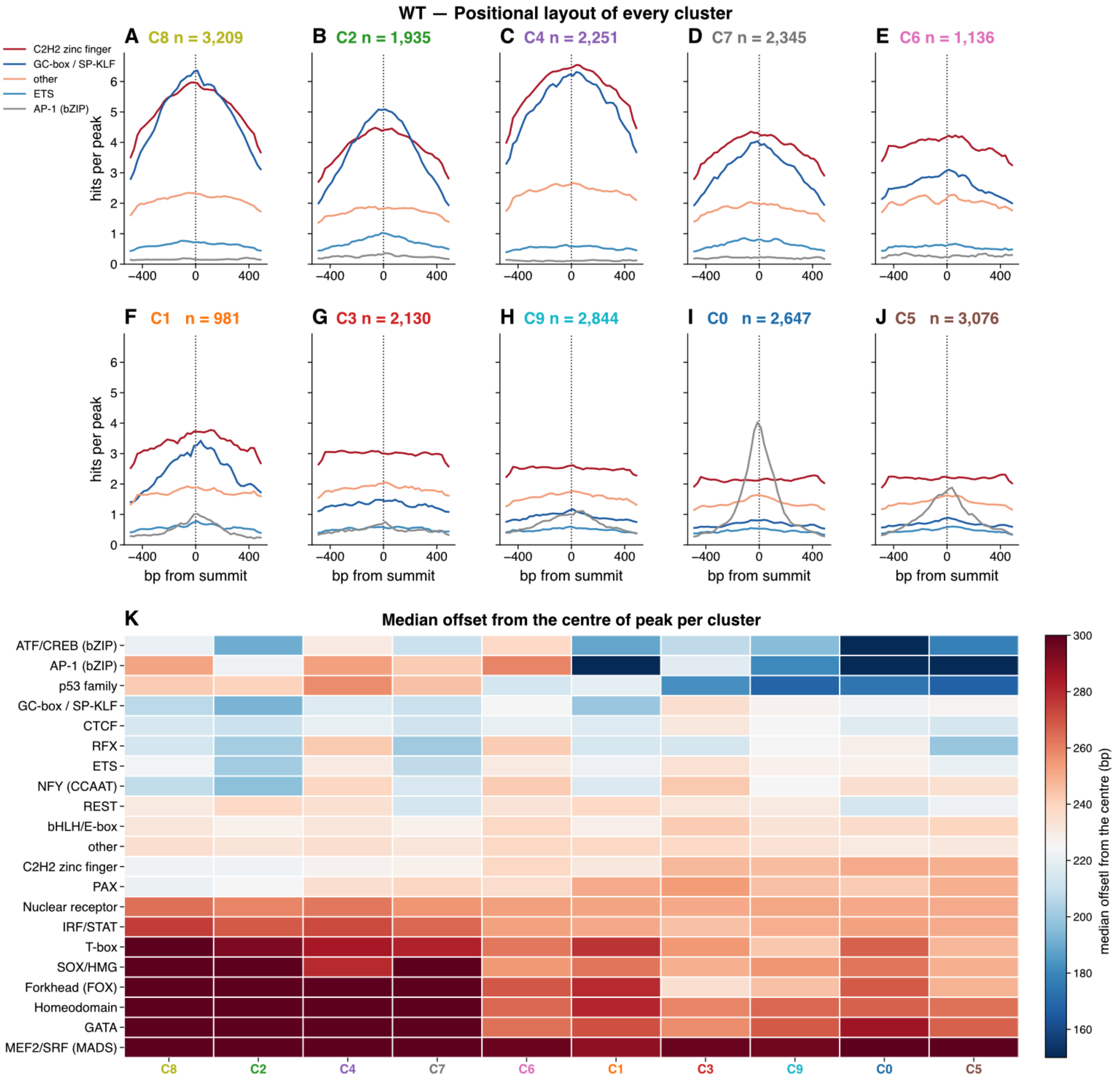


**Supplementary Figure 8.** Positional distribution of transcription factor binding motifs across predicted WT clusters. **(A–J)** Metaplots showing motif hits per peak, grouped by TF family, across ±500 bp from the peak summit for each predicted cluster. **(K)** Heatmap showing the median offset from the peak centre for each TF family across clusters.
